# Brain-wide single neuron reconstruction reveals morphological diversity in molecularly defined striatal, thalamic, cortical and claustral neuron types

**DOI:** 10.1101/675280

**Authors:** Hanchuan Peng, Peng Xie, Lijuan Liu, Xiuli Kuang, Yimin Wang, Lei Qu, Hui Gong, Shengdian Jiang, Anan Li, Zongcai Ruan, Liya Ding, Chao Chen, Mengya Chen, Tanya L. Daigle, Zhangcan Ding, Yanjun Duan, Aaron Feiner, Ping He, Chris Hill, Karla E. Hirokawa, Guodong Hong, Lei Huang, Sara Kebede, Hsien-Chi Kuo, Rachael Larsen, Phil Lesnar, Longfei Li, Qi Li, Xiangning Li, Yaoyao Li, Yuanyuan Li, An Liu, Donghuan Lu, Stephanie Mok, Lydia Ng, Thuc Nghi Nguyen, Qiang Ouyang, Jintao Pan, Elise Shen, Yuanyuan Song, Susan M. Sunkin, Bosiljka Tasic, Matthew B. Veldman, Wayne Wakeman, Wan Wan, Peng Wang, Quanxin Wang, Tao Wang, Yaping Wang, Feng Xiong, Wei Xiong, Wenjie Xu, Zizhen Yao, Min Ye, Lulu Yin, Yang Yu, Jia Yuan, Jing Yuan, Zhixi Yun, Shaoqun Zeng, Shichen Zhang, Sujun Zhao, Zijun Zhao, Zhi Zhou, Z. Josh Huang, Luke Esposito, Michael J. Hawrylycz, Staci A. Sorensen, X. William Yang, Yefeng Zheng, Zhongze Gu, Wei Xie, Christof Koch, Qingming Luo, Julie A. Harris, Yun Wang, Hongkui Zeng

**Affiliations:** Allen Institute for Brain Science, Seattle, WA 98109, USA; SEU-ALLEN Joint Center, Institute for Brain and Intelligence, Southeast University, Nanjing, Jiangsu, China; School of Optometry and Ophthalmology, Wenzhou Medical University, Wenzhou, Zhejiang, China; School of Computer Engineering and Science, Shanghai University, Shanghai, China; Key Laboratory of Intelligent Computation & Signal Processing, Ministry of Education, Anhui University, Hefei, Anhui, China; Britton Chance Center for Biomedical Photonics, Wuhan National Laboratory for Optoelectronics, MoE Key Laboratory for Biomedical Photonics, Huazhong University of Science and Technology, Wuhan, Hubei, China; HUST-Suzhou Institute for Brainsmatics, JITRI Institute for Brainsmatics, Suzhou, Jiangsu, China; Tencent Jarvis Lab, Shenzhen, Guangdong, China; Center for Neurobehavioral Genetics, Jane and Terry Semel Institute for Neuroscience and Human Behavior; Department of Psychiatry and Biobehavioral Sciences, David Geffen School of Medicine, University of California, Los Angeles, Los Angeles, CA, USA; Cold Spring Harbor Laboratory, Cold Spring Harbor, NY, USA; School of Biomedical Engineering, Hainan University, Haikou, Hainan, China

## Abstract

Ever since the seminal findings of Ramon y Cajal, dendritic and axonal morphology has been recognized as a defining feature of neuronal types. Yet our knowledge concerning the diversity of neuronal morphologies, in particular distal axonal projection patterns, is extremely limited. To systematically obtain single neuron full morphology on a brain-wide scale, we established a platform with five major components: sparse labeling, whole-brain imaging, reconstruction, registration, and classification. We achieved sparse, robust and consistent fluorescent labeling of a wide range of neuronal types by combining transgenic or viral Cre delivery with novel transgenic reporter lines. We acquired high-resolution whole-brain fluorescent images from a large set of sparsely labeled brains using fluorescence micro-optical sectioning tomography (fMOST). We developed a set of software tools for efficient large-volume image data processing, registration to the Allen Mouse Brain Common Coordinate Framework (CCF), and computer-assisted morphological reconstruction. We reconstructed and analyzed the complete morphologies of 1,708 neurons from the striatum, thalamus, cortex and claustrum. Finally, we classified these cells into multiple morphological and projection types and identified a set of region-specific organizational rules of long-range axonal projections at the single cell level. Specifically, different neuron types from different regions follow highly distinct rules in convergent or divergent projection, feedforward or feedback axon termination patterns, and between-cell homogeneity or heterogeneity. Major molecularly defined classes or types of neurons have correspondingly distinct morphological and projection patterns, however, we also identify further remarkably extensive morphological and projection diversity at more fine-grained levels within the major types that cannot presently be accounted for by preexisting transcriptomic subtypes. These insights reinforce the importance of full morphological characterization of brain cell types and suggest a plethora of ways different cell types and individual neurons may contribute to the function of their respective circuits.

## INTRODUCTION

Understanding the taxonomic organization of cell types in the brain will yield insight into its fundamental components and landscape, providing an atlas much like the periodic table of elements in chemistry and the taxonomy of living species in biology. As neurons exhibit extraordinary diversity across molecular, morphological, physiological, and connectional features, a complete and accurate classification and creation of a cell type atlas needs to consider and integrate these distinct cellular properties ^1^. Recent advances in high-throughput single cell RNA-sequencing has enabled the systematic classification of cell types at the transcriptomic level ^2–6^. This approach captures major cell types with known anatomical and functional properties, but also reveals many potential new cell types. Systematic classification of cortical neurons using a combination of local morphological and electrical properties has also been achieved ^7–9^, and new technologies have been developed to correlate morpho-electrical and transcriptomic cell types ^10, 11^. Although dendritic and long-range axonal morphologies have long been held as the central defining feature of neuronal types ^12^, so far very few tools are available and little systematic effort has been made to classify individual single neurons using brain-wide axonal projection patterns alone or with other cellular properties.

Brain-wide inter-areal connectivity has been mapped extensively using injections of anterograde and retrograde tracers to label populations of projection neurons ^13–18^. However, it remains largely unknown how population-level projection patterns are reflected at the single neuron level. Triple retrograde tracing studies suggest that individual neurons within a brain region often have heterogeneous axonal projection patterns ^19–21^. Thus, characterizing single neuron axonal projections through reconstruction of complete morphologies not only provides critical information related to classifying cell types, but also how neural signals are organized and transmitted to their target regions.

Despite its importance, data on single neuron axonal morphologies are currently lacking for most projection neuron types in mammals, in large part because axons often cover large distances and are severed in *ex vivo* brain slices. Previous efforts have been made in rodents to fully label single neurons with small molecules or fluorescent proteins through *in vivo* whole-cell patching, *in vivo* electroporation ^22–24^, sparse transgenic labeling ^25^, or sparse viral labeling with sindbis virus ^26–28^ or adeno-associated virus (AAV) ^29–31^. Conventionally this is followed by serial sectioning, imaging of each section, and manual reconstruction of the labeled neurons across many consecutive sections. Although relatively few such studies exist due to the labor-intensive process, they reveal critical features of specific projection neuron types that likely have important functional implications ^22, 24, 32–39^. The recent development of high-throughput and high-resolution fluorescent imaging platforms, such as fMOST ^40^ and MouseLight ^29^, coupled with more efficient sparse viral labeling strategies, now enable the potential for large-scale generation of neuronal morphology datasets. These studies also revealed a need for further improvements in tools for generating very sparse and strong labeling of single neurons at a brain-wide scale, as well as computational tools to expedite the laborious reconstruction process.

As part of the BRAIN Initiative Cell Census Network (BICCN) efforts to characterize brain cell types across multiple modalities, we have established a pipeline to label, image, reconstruct and classify single neurons at a brain-wide scale using complete morphology data. As a foundational step toward this goal, we report here the largest set of single neuron reconstructions to this date, 1,708 neurons from striatum, thalamus, cortex, claustrum and other brain regions in mice, as a multi-site collaborative effort. These neurons are labeled by cell class or type selective Cre driver lines, enabling correlation of their morphologies and projection patterns with their molecular identities. We also provide a corresponding set of single-cell RNA-seq data from retrogradely labeled neurons (i.e. Retro-seq) to corroborate our findings. Overall, our study reveals substantial morphological and projection diversity of individual neurons; this diversity is governed by underlying rules that are region- and cell type-specific. The results demonstrate that full morphology reconstruction is an essential component of cell type characterization, highly complementary to other molecular, anatomical and physiological approaches in order to gain a full understanding of cell type diversity. To facilitate and encourage further work in this key area all imaging data and computational reconstruction tools are made publicly available. Our ultimate goal is to enable and encourage a community-based effort to generate a sufficiently large set of full morphology reconstructions, potentially tens to hundreds of thousands, to facilitate cell type classification, comparison with molecular, physiological and other cellular properties and understanding the functional implications of single neurons.

## RESULTS

### Sparse, robust and consistent neuronal labeling

Full reconstruction of a single neuron requires sparse, robust and consistent neuronal labeling. Previous genetic approaches applied to produce sparse or single neuron labeling, including viral delivery (*e.g.*, using sindbis virus or AAV) and *in vivo* electroporation ^22–24, 26–31, 37^, often resulted in substantial cell-to-cell and animal-to-animal variations, and were usually restricted to few brain regions. In rare cases, extremely sparse labeling was obtained utilizing the low recombination efficiency of CreER driver lines and a sensitive, alkaline phosphatase-based histochemical reporter ^25, 41^. To achieve more efficient, widespread, consistently sparse yet strong labeling, we utilized TIGRE2.0 transgenic reporter lines that exhibit viral-like transgene expression levels ^42, 43^, coupling them with Cre expression from either driver lines or viral delivery. We employed two general approaches.

The first was to use the GFP expressing Ai139 or Ai140 TIGRE2.0 reporter line in conjunction with sparse Cre-mediated recombination (Fig. 1a). We used CreERT2 driver lines and titrated the level of CreERT2-mediated recombination using low-dose tamoxifen induction (**Supplementary Table 1**). We found optimal tamoxifen doses for sparse labeling in each case using serial two photon tomography (STPT) to quickly screen for brain-wide transgene expression ^16^ (Extended Data Fig. 1a-c). We also tested a dual reporter combination with TIGRE2.0 (Ai140) and TIGRE1.0 (Ai82) ^44^ reporter lines, using the tTA2 from the single TIGRE2.0 allele to drive two copies of the TRE promoter driven GFP expression cassettes (Fig. 1a, “optional” cross). We found this strategy generated an even higher level of GFP expression than Ai139 or Ai140 alone, well suited for fMOST imaging (see below).

**Figure 1.**
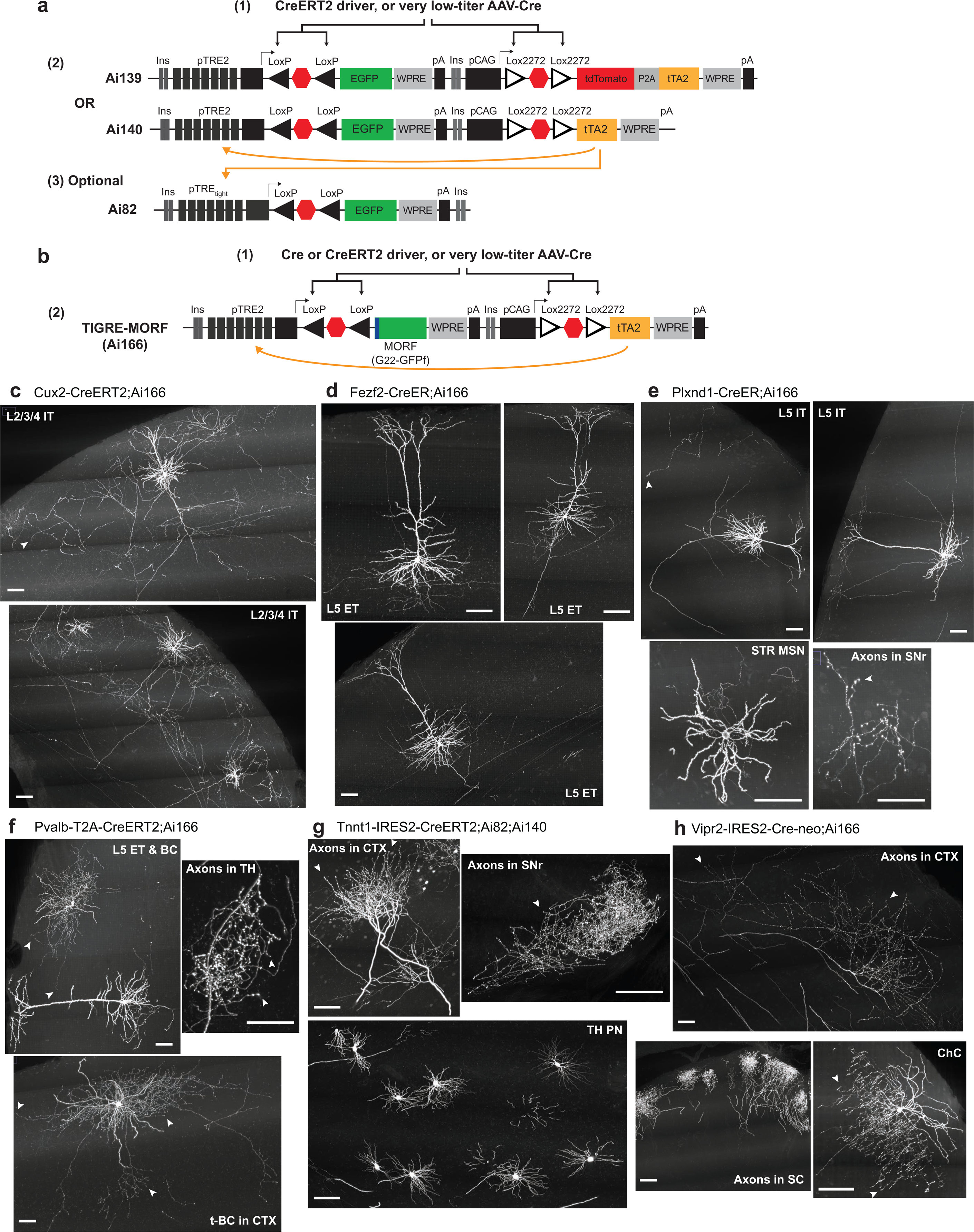
Sparse, robust and consistent labeling and visualization of the dendritic and axonal arborizations of a wide range of neuronal types. **a**, Schematic diagram showing the combination of CreERT2 transgenic driver line or Cre-expressing AAV (1) with the GFP-expressing TIGRE2.0 reporter line Ai139 or Ai140 (2). Very low dose tamoxifen induction of CreERT2 or very low-titer AAV-Cre delivery results in activation of the reporter in a spatially sparse manner. Transgenic reporter expression of GFP is robust and consistent across different cells. An optional addition is to cross in the GFP-expressing TIGRE1.0 reporter line Ai82 (3), so that the tTA2 from Ai139 or Ai140 will activate the expression of GFP from two alleles – Ai139/Ai140 and Ai82, further increasing the level of GFP within Cre+ cells. **b**, Schematic diagram showing the combination of Cre or CreERT2 transgenic driver line or Cre-expressing AAV (1) with the GFP-expressing sparse reporter line TIGRE-MORF/Ai166 (2). Due to the intrinsic sparse expression of MORF (G_22_-GFPf), some conventional Cre lines, moderate doses of tamoxifen induction of CreERT2, or moderate titers of AAV-Cre delivery can result in very sparse labeling. **c**, Cortical L2/3/4 IT neurons and their extensive local axon collaterals clearly labeled in a Cux2-CreERT2;Ai166 brain. **d**, Cortical L5 ET neurons and their sparse local axon collaterals seen in a Fezf2-CreER;Ai166 brain. **e**, Cortical L5 IT neurons and their local axon collaterals seen in a Plxnd1-CreER;Ai166 brain. Striatal medium spiny neurons (STR MSN) are also sparsely labeled, and their individual axons are clearly seen in substantia nigra (SN). **f**, Cortical inhibitory basket cells (BC) and translaminar basket cells (t-BC), as well as L5 ET excitatory neurons, seen in a Pvalb-T2A-CreERT2;Ai166 brain. The L5 ET neurons form driving-type axon clusters with large boutons in the thalamus (TH). **g**, Thalamic projection neurons (TH PN) with their dense axon terminal clusters in cortex seen in a Tnnt1-IRES2-CreERT2;Ai82;Ai140 brain. Some STR MSNs are also labeled and they form intense axon clusters in SN. **h**, In a Vipr2-IRES2-Cre-neo;Ai166 brain, axon clusters from projection neurons in visual thalamic nuclei are seen in CTX, axon clusters likely from retinal ganglion cells are seen in superior colliculus (SC), and a cortical chandelier cell (ChC) is fully labeled with its characteristic axonal branches. Images shown in c-h are 100-µm maximum intensity projection (MIP) images (*i.e.*, projected from 100 consecutive 1-µm image planes). Arrowheads indicate observed terminal boutons at the end of the axon segments. Tamoxifen doses are shown in **Supplementary Table 1**. Scale bars, 100 µm.

The second approach was to use a new TIGRE2.0 Cre reporter line: TIGRE-MORF (also called Ai166) ^43^, which is sparsely activated in conjunction with Cre delivery (Fig. 1b). TIGRE-MORF/Ai166 expresses the MORF gene, which is composed of a farnesylated EGFP (GFPf) preceded by a mononucleotide repeat of 22 guanines (G_22_-GFPf). The GFPf transgene is not translated at the baseline due to the out-of-frame G_22_ repeat relative to the open reading frame of GFPf, which lacks its own translation start codon. However, during DNA replication or repair, rare events of stochastic frameshift of the mononucleotide repeat result in correction of the translation frame (*e.g.*, G_22_ to G_21_) and produce expression of the GFPf protein in a small subset of cells. TIGRE-MORF/Ai166 (below simplified as Ai166) exhibits a labeling frequency of 1-5% when crossed to different Cre driver mouse lines ^43^. Even with this frequency, we found that combining Ai166 with many Cre driver lines densely expressing the Cre transgene did not produce sufficient sparsity to readily untangle the axonal ramifications, whereas combining it with Cre lines that are already relatively sparse to begin with, or with CreERT2 lines with intermediate dosing level of tamoxifen (**Supplementary Table 1**), leads to extremely sparse labeling well suited for reconstruction of elaborate axonal arborizations of many neuronal types (Extended Data Fig. 1d-j). The use of membrane associated GFPf also enabled robust labeling of very thin axon fibers. Leaky background expression of GFP reported in other TIGRE2.0 lines ^42^ is not present in Ai166 mice due to the strict dependency of translational frameshift for the expression of GFPf reporter, making TIGRE-MORF/Ai166 an ideal reporter line for sparse and strong labeling of various neuronal types across the brain.

### High quality imaging data reveals diverse neuronal morphologies

For this study, we generated 53 high-quality (*i.e.*, strong, even and not-too-dense labeling) fMOST-imaged brain datasets with sparsely labeled cells in cortical, thalamic, claustral, and striatal regions, and for cholinergic, noradrenergic and serotonergic neuronal types (Fig. 1c-h, Extended Data Fig. 2, **Supplementary Table 1**). Critically, our approach can be extended to any cell type for which appropriate Cre-dependent labeling methods are available. In the cortex, we imaged from different excitatory projection classes using selective CreERT2 driver lines ^4, 42, 45^. For example, Cux2-CreERT2;Ai166 labeled the cortical layer (L) 2/3/4 intratelencephalic (IT) subclasses of excitatory neurons (Fig. 1c). Plxnd1-CreER;Ai166 labeled cortical L2/3 and L5 IT subclasses, as well as striatal medium spiny neurons (MSN, Fig. 1e). Fezf2-CreER;Ai166 labeled cortical L5 extratelencephalic (ET, also known as pyramidal tract, PT) subclass (Fig. 1d). Tle4-CreER;Ai166 labeled L6 corticothalamic (CT) subclass (Extended Data Fig. 2a). Nxph4-T2A-CreERT2;Ai166 labeled cortical L6b subplate neurons (Extended Data Fig. 2b). In the cortex, we also labeled and imaged *Pvalb*+ cells, which includes a subclass of inhibitory interneurons, *e.g.*, basket cells (BC), and a subset of L5 ET excitatory neurons, using Pvalb-T2A-CreERT2;Ai166 (Fig. 1f) and *Sst*+ interneurons using Sst-Cre;Ai166 (Extended Data Fig. 2c). In the thalamus, we used Tnnt1-IRES2-CreERT2;Ai140;Ai82 and Vipr2-IRES2-Cre-neo;Ai166 to label excitatory projection neurons as well as striatal MSNs (Fig. 1g,h). Vipr2-IRES2-Cre-neo;Ai166 also labeled axons consistent with projections from retinal ganglion cells ^46^, as well as cortical chandelier cells (ChC) (Fig. 1h). Gnb4-IRES2-CreERT2;Ai140;Ai82 labeled the *Car3*+ IT subclass of L6 excitatory neurons in cortex and claustrum (CLA, Extended Data Fig. 2d). Cell types containing neuromodulators were also labeled with selective Cre driver lines ^47^, including noradrenergic neurons in the locus ceruleus (LC) using Dbh-Cre_KH212;Ai166 (Extended Data Fig. 2e) and serotonergic neurons in the dorsal raphe (DR) and other brainstem regions using Slc6a4-CreERT2;Ai166 (Extended Data Fig. 2f).

**Figure 2.**
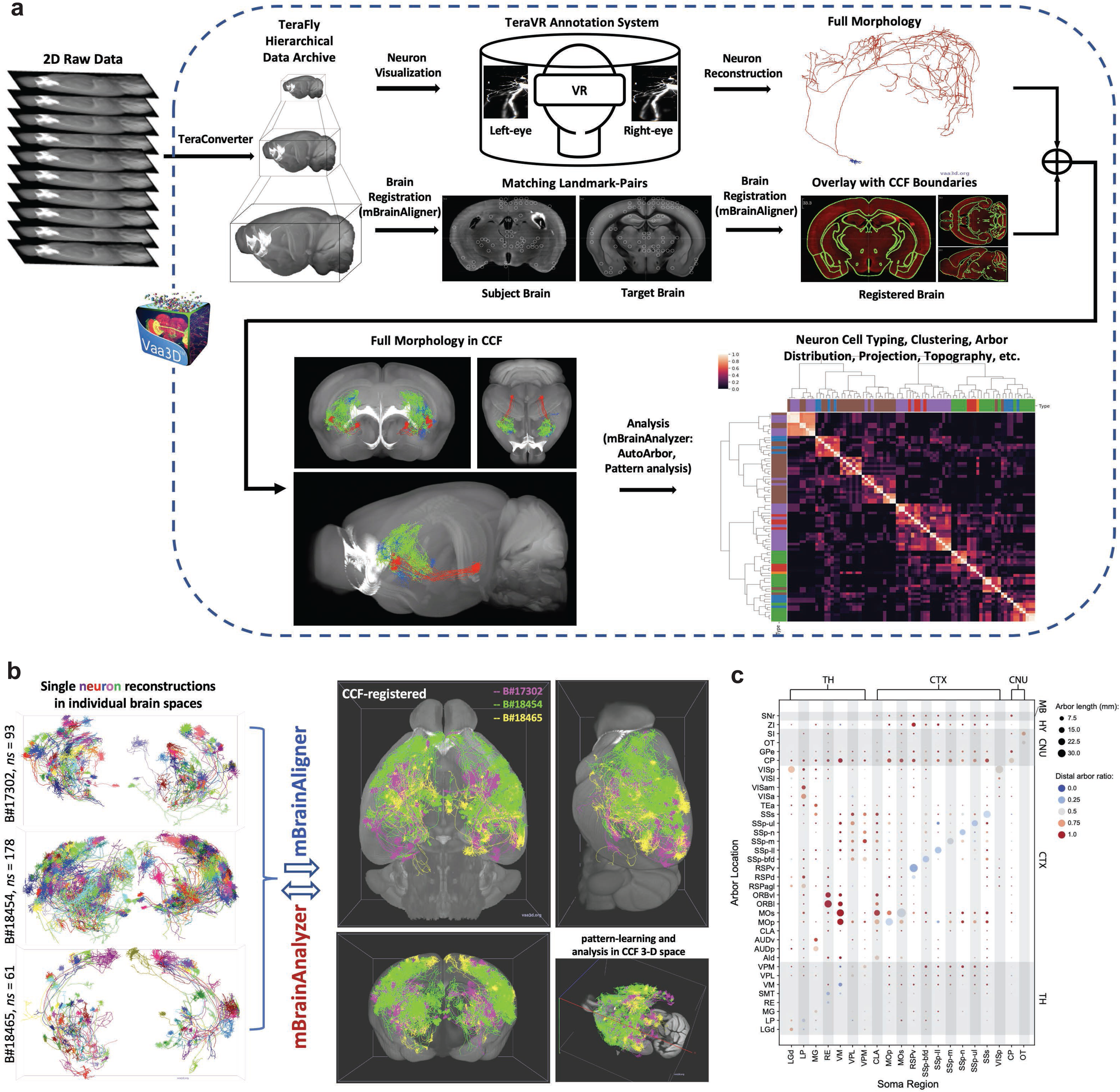
Platform and workflow of the brain-wide full morphology reconstruction, registration and analysis pipeline. **a**, The workflow of neuron visualization, reconstruction, mapping to Common Coordinate Framework (CCF) and analysis. A complete fMOST image dataset is first converted to TeraFly file format by TeraConverter, the data formatting tool in TeraFly. Then annotators work in the TeraVR annotation system to reconstruct the full morphology of each neuron. In parallel, the whole brain image dataset is registered to CCF using mBrainAligner, which uses both RLM (Reliable-Landmark-Matching) and LQM (Little-Quick-Warp) modules in brain alignment. Following registration of the image dataset to CCFv3, all the reconstructed morphologies from the same brain are also registered for subsequent visualization and quantitative analysis. **b**, Demonstration of single neuron reconstructions shown in varying colors from individual mouse brains registered to the CCF space by mBrainAligner, which allows integrated analysis by the morphology analysis toolbox, mBrainAnalyzer. **c**, Overview of projection patterns. Cells are grouped by curated soma locations, first by major brain areas (e.g. CTX: cortex, TH: thalamus, CNU: Cerebral nuclei) and then by refined areas (e.g. LGd nucleus of the thalamus). Sizes of dots represent group-average of arbor length. Colors represent the ratio of dendritic arbors (red: low; blue: high). Note that some minor “projections” indicated by tiny dots may be false positives due to passing fibers or not completely precise registration.

It is apparent that these neurons display a remarkable array of dendritic and axonal morphologies. Specifically, in these sparsely labeled brains, cortical IT and ET neurons not only have primary long-range projections but also local axonal branches that are well segregated and clearly identifiable, enabling truly complete reconstruction of the entire local and long-range, cortical and subcortical axonal arborization (Fig. 1c-e). L5 ET neurons form the ‘driving’ type of synapses in the thalamus ^48, 49^, which have enlarged and intensely fluorescent boutons (Fig. 1f). L6b subplate neurons extend their local axon collaterals upwards into layer 1 (Extended Data Fig. 2b). The axons of thalamic projection neurons form either dense or dispersed clusters in the cortex (Fig. 1g-h). On the other hand, claustral, noradrenergic and serotonergic neurons have widely dispersed, thin axons that are nonetheless well labeled (Extended Data Fig. 2d-f). One can also clearly see individual axons in the substantia nigra from striatal medium spiny neurons (Fig. 1e,g), individual axon terminal clusters in the superficial layers of the superior colliculus likely coming from retinal ganglion cells (Fig. 1h), as well as the dense and fine local axonal branches of a variety of cortical and striatal interneurons (*e.g.*, basket cells, chandelier cells, and Martinotti cells) (Fig. 1f,h and Extended Data Fig. 2c).

Of note, sparsely labeled neurons were frequently observed in other regions of the brain for all of these crosses but are not described in detail here. Each of these brains contains ∼100-1,000 labeled neurons (**Supplementary Table 1**). Thus, tens of thousands of neurons could be reconstructed from these and newly generated datasets in the coming years. The whole brain image series are publicly available through the BICCN web portal (https://biccn.org/) as a unique resource for the community.

### Pipeline for image data processing, morphology reconstruction and registration

We acquired whole brain images with sufficient resolution (∼0.3 x 0.3 x 1 μm XYZ) for reconstructing fine-caliber axons using fluorescence micro-optical sectioning tomography (fMOST), a high-throughput, high-resolution, brain-wide fluorescent imaging platform ^40^. To handle the large imaging datasets generated, we established a standardized image data processing and informatics workflow (Fig. 2a) for efficient whole brain morphology reconstruction utilizing Vaa3D, an open-source, cross-platform visualization and analysis system ^50, 51^. Each fMOST dataset is first converted to a multi-level navigable dataset using the Vaa3D-TeraFly program ^52^, which allows smooth handling of terabyte-scale datasets. Neuron visualization and reconstruction is then carried out on the TeraFly files. A series of tools, especially those based on the “Virtual Finger” method ^53^, were developed within Vaa3D to facilitate semi-automated and manual reconstruction. Further, a virtual reality (VR) environment created within Vaa3D, TeraVR, significantly enhances a user’s ability to see the 3D relationships among intertwined axonal segments, improving precision and efficiency of reconstruction ^54^. After quality control (QC) and manual correction, we used Vaa3D’s deformable model to automatically fit the tracing to the center of fluorescent signals. The final reconstructed morphology was completed as a single tree without breaks, loops, or trifurcations. All these data processing, reconstruction, and workflow control processes were managed using a newly designed software system for massive scale data production (Jiang et al, manuscript in preparation).

In parallel, each fMOST dataset was registered to the 3D Allen mouse brain Common Coordinate Framework (CCFv3, http://atlas.brain-map.org/) ^55^, using a newly developed mBrainAligner program (Methods; Qu et al, manuscript in preparation) specifically designed for fMOST datasets to handle the challenges of brain shrinkage and deformation related to modality-specific technical protocols (Fig. 2b, Extended Data Fig. 3). Following registration of the whole-brain image dataset, all individual neuron reconstructions were also registered to the CCFv3 using the source brain’s transformation parameters (Fig. 2b). Registration to CCFv3 enables digital anatomical delineation and spatial quantification of each reconstructed morphology and its compartments (*e.g.*, soma, dendrites, axon arbors). Since neurons are reconstructed from different brains, co-registration to the CCFv3 allows them to be compared and analyzed using a unified framework, mBrainAnalyzer (Fig. 2b, Extended Data Fig. 3), which automatically detects the arbors of each neuron followed by mapping of these dendritic and axonal arbors onto the standardized CCFv3 space. Using these tools and following this workflow, we reconstructed the full 3D morphology of 1,708 neurons from cortex, claustrum, thalamus, striatum and other regions (**Supplementary Table 2**). Quantification of all detected axon arbors, shown with respect to the anatomical locations of their somas, provides an informative global view of the brain-wide projection patterns for these single neurons (Fig. 2c). Detailed analysis of this brain-wide projection arbor map to understand the morphological diversity of cell types sheds light on how to perform more comprehensive analysis of cell types with finer resolution and precision.

**Figure 3.**
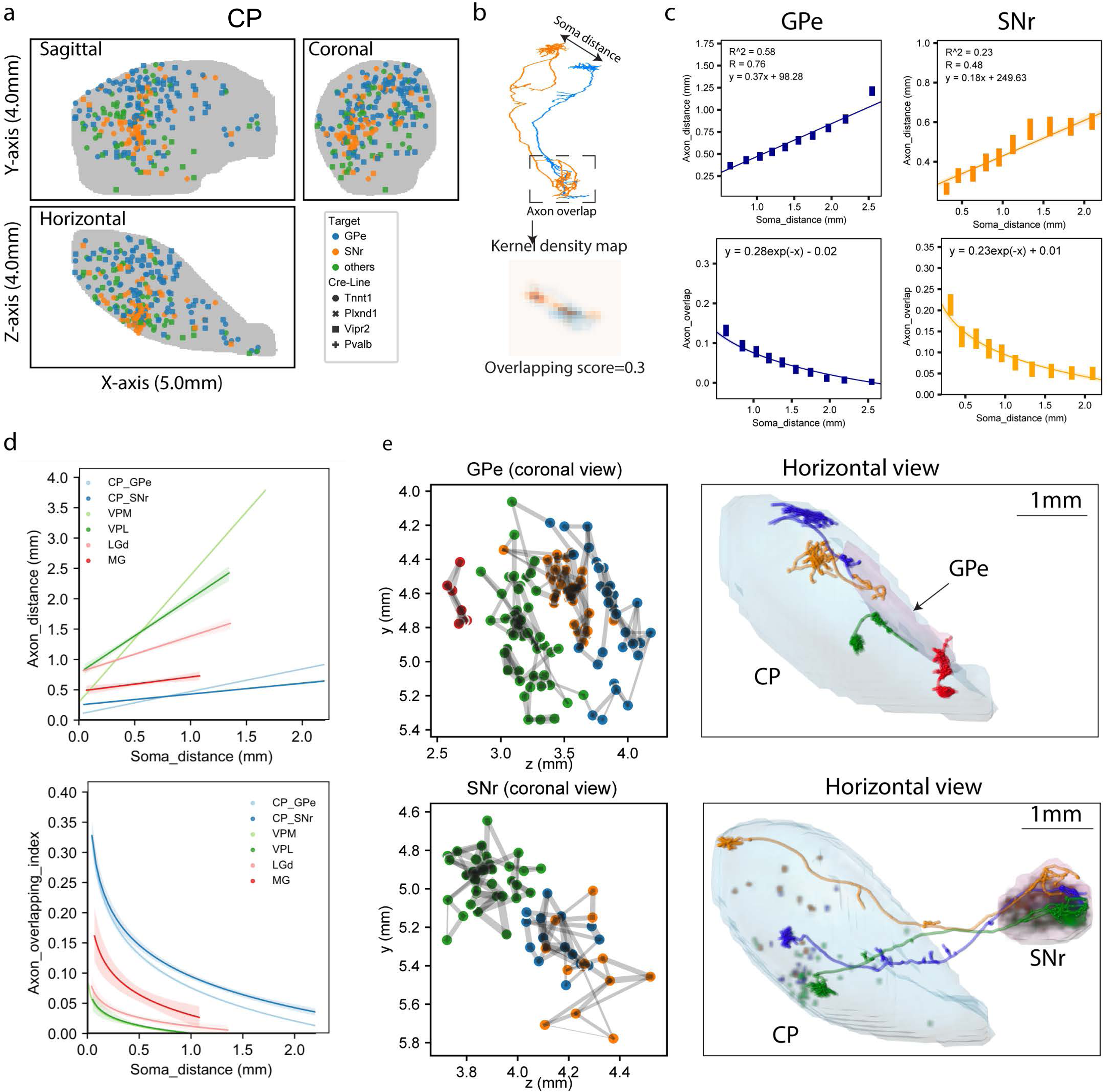
Striatal neuron morphology analysis. **a**, Coronal, sagittal and horizontal views of soma distribution of CP neurons (X-axis: anterior to posterior, Y-axis: dorsal to ventral, Z-axis: medial to lateral. The same axis labels were used throughout the paper). **b,** Overlapping score of axons is calculated by estimating the kernel density map of individual axon arbors and the density-weighted average of overlapping areas for each arbor pair. **c,** Regression of distance between arbor centers (top panels) or overlapping score (bottom panels) by soma distance. Linear and negative exponential models are used for distance and overlapping score, respectively. **d,** Comparison of arbor convergence across cell types. Regression curves generated by the same approach as in c. Colors represent cell types. Light-shaded bands represent 95% confidence intervals. **e,** Clustering of axon overlapping by Louvain algorithm. Left panels, coronal views of axon arbor locations colored by clusters. Width of grey lines represents overlapping scores between arbor pairs. Right panels, representative single neurons to illustrate topography of CP neuron projections. Cells are colored by cluster identities.

We established a stringent QC process that includes ensuring the completeness of reconstructed morphologies (Extended Data Fig. 4). A conventional way to assess the completeness of axon labeling and reconstruction is whether an axon ends at a bouton (indicated by an enlargement with more intense signal (see arrowheads in Fig. 1) or gradually tapers off, the former suggesting a complete labeling ^36^. We implemented this assessment in our reconstruction refinement process to identify potential inaccuracies (Extended Data Fig. 4c-e). In our final QC-passed reconstructions we found that the ratio between terminal axon branches with and without a terminal bouton was about 10:1, indicating a high degree of completeness of our reconstructed morphologies.

**Figure 4.**
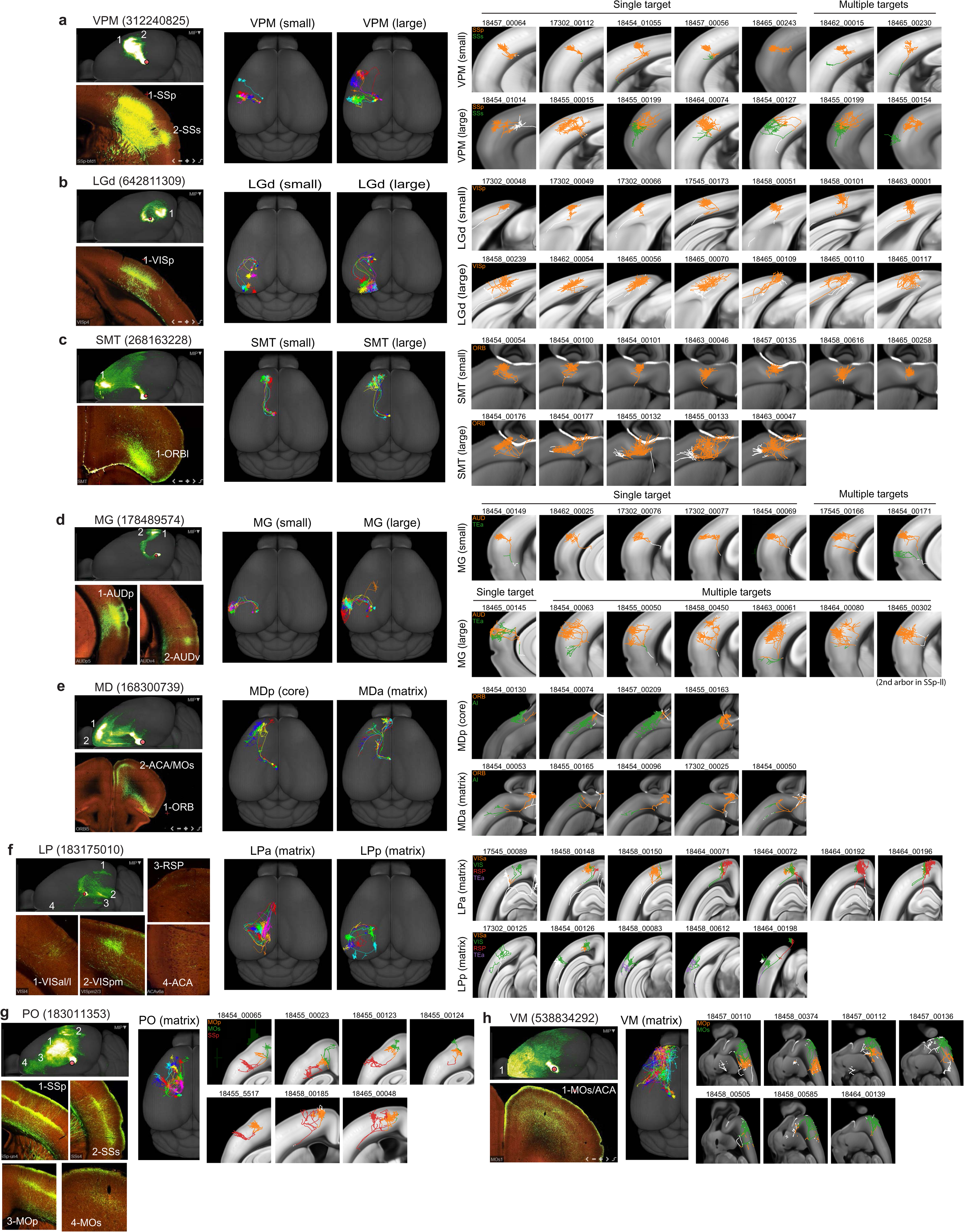
Long-range projection patterns of individual thalamic neurons in comparison with mesoscale population-level projections. **a-h,** Axonal morphologies and projections of reconstructed single neurons compared with population projection patterns for nucleus VPM (a), LGd (b), SMT (c), MG (d), MD (e), LP (f), PO (g) and VM (h). For each nucleus, left panels, representative mesoscale experiments shown in a maximum projection whole-brain top-down view and individual higher-power images showing axon termination patterns in major target regions; middle panels, all example single neurons shown together in a maximum projection whole-brain top-down view; right panels, each neuron is shown in a chosen plane to best capture the perpendicular (to pial surface) orientation of the main axon arbor with superimposed maximum projection view of the neuron’s axon arbors. The chosen plane can be coronal (for a, b, d), horizontal (for c, e), sagittal (for h) or tilted (for f, g), based on the main cortical target region. Different cortical target regions are indicated by different colors. Small, small axon arbors. Large, large axon arbors. MDa and MDp, or LPa and LPp are the anterior and posterior parts of MD or LP respectively.

We calculated a set of morphological features (Methods) for every neuron, focused on soma location, dendrite and axonal properties. We also used a graph-partition approach to automatically detect the major domains of arborization for each neuron (Methods). Morphological features such as length, depth, area, etc., at the whole neuron level were also computed for each arbor-domain for analysis. In the following sections, we use a combination of these metrics to classify and analyze single neurons in several major brain divisions.

### Striatal medium spiny neurons exhibit converging projections while retaining topography

We reconstructed 305 neurons in the dorsal striatum (caudate putamen, CP) from 4 Cre driver lines: Tnnt1, Plxnd1, Vipr2 and Pvalb (Fig. 3a, **Supplementary Table 2**). These neurons can be divided into 3 groups based on their projection targets: (1) those with main axon projections terminating in the external segment of the globus pallidus (GPe, n=179 cells), (2) those terminating in the reticular part of the substantia nigra (SNr, n=96 cells) and (3) those terminating within striatum itself (others, n=30 cells). These groups of neurons are distributed throughout CP and largely intermingled. Axonal projections from groups 1 and 2 correspond to the two well-known types of striatal medium spiny neurons, dopamine receptor D1 (*Drd1*) neurons projecting to SNr and dopamine receptor D2 (*Drd2*) neurons projecting to GPe ^56^.

Individual striatal neurons projecting to GPe or SNr exhibit a simple point-to-point morphology: GPe-projecting neurons have one major axon arbor, targeting GPe; SNr-projecting neurons have one major axon arbor targeting SNr. The GPe type has more elaborate axon arborization near the soma. Sholl analysis shows that the number of local crossings (<1 mm to soma) of the GPe type is 2.8 times of that of the SNr type. The radius of local axons of the GPe type is also 2.5 times as large. Most SNr-projecting neurons send minor branches targeting the internal segment of globus pallidus (GPi) or GPe.

The dominant feature of both types of striatal neurons is convergent projection within the main target region, GPe or SNr, consistent with the ∼20-fold smaller sizes/volumes of these regions compared to the dorsal striatum (Fig. 3b,c). We find that both the center-to-center distance and the degree of overlap of axon arbors between each pair of neurons are proportional to that pair’s soma-to-soma distance (Fig. 3c), indicating a regular spatial organization of these neurons’ axon projections. The axon arbor distances between striatal neurons within the same type are substantially smaller and their overlapping scores are substantially greater in comparison to those of neurons from each of the various thalamic nuclei (see next section) (Fig. 3d). Furthermore, axon arbors in GPe or SNr can be grouped into domains based on the degree of overlap; these domains are arranged topographically and correspond to the topographic localization of the somas in striatum (Fig. 3e).

### Thalamic neurons exhibit diverse and nucleus-specific thalamocortical projection patterns

We reconstructed 735 thalamic neurons from 3 Cre lines: Tnnt1, Vipr2 and Pvalb (**Supplementary Table 2**). Those (n=17 neurons) from Pvalb-Cre are mostly GABAergic neurons and in the reticular nucleus (RT) projecting back to other thalamic nuclei. Tnnt1 and Vipr2 Cre lines contained mostly thalamocortical projection neurons from many (but not all) nuclei, including sensory-motor relay nuclei (n=658 cells) and higher-order or associational nuclei (n=60 cells). In this dataset, the reconstructed cells covered 22 of the 44 thalamic regions in CCFv3, which can be broadly divided into two major groups ^49, 57, 58^: (1) primary sensory or motor relay nuclei, also known as “core” or “driver” nuclei, which include the ventral posteromedial nucleus (VPM), ventral posterolateral nucleus (VPL), ventral posteromedial nucleus parvicellular part (VPMpc), ventral posterolateral nucleus parvicellular part (VPLpc), dorsal part of the lateral geniculate complex (LGd), medial geniculate complex (MG) and ventral anterior-lateral complex (VAL), as well as anterior and medial nuclei such as anteromedial nucleus (AM), submedial nucleus (SMT) and the posterior part of the mediodorsal nucleus (MD) (Extended Data Fig. 5); (2) associational or higher-order thalamic nuclei, also known as “matrix” or “modulatory” nuclei, which include the lateral posterior nucleus (LP), posterior complex (PO), lateral dorsal nucleus (LD), ventral medial nucleus (VM), nucleus of reuniens (RE), central medial nucleus (CM), interanterodorsal nucleus (IAD), anterior part of MD, and paraventricular nucleus (PVT) (Extended Data Fig. 6). In general, “core-type” thalamocortical projections target one or a small number of cortical regions and predominantly terminate in L4. In contrast, the “matrix-type” thalamocortical projections generally target multiple cortical areas with dense axonal terminals in L1.

**Figure 5.**
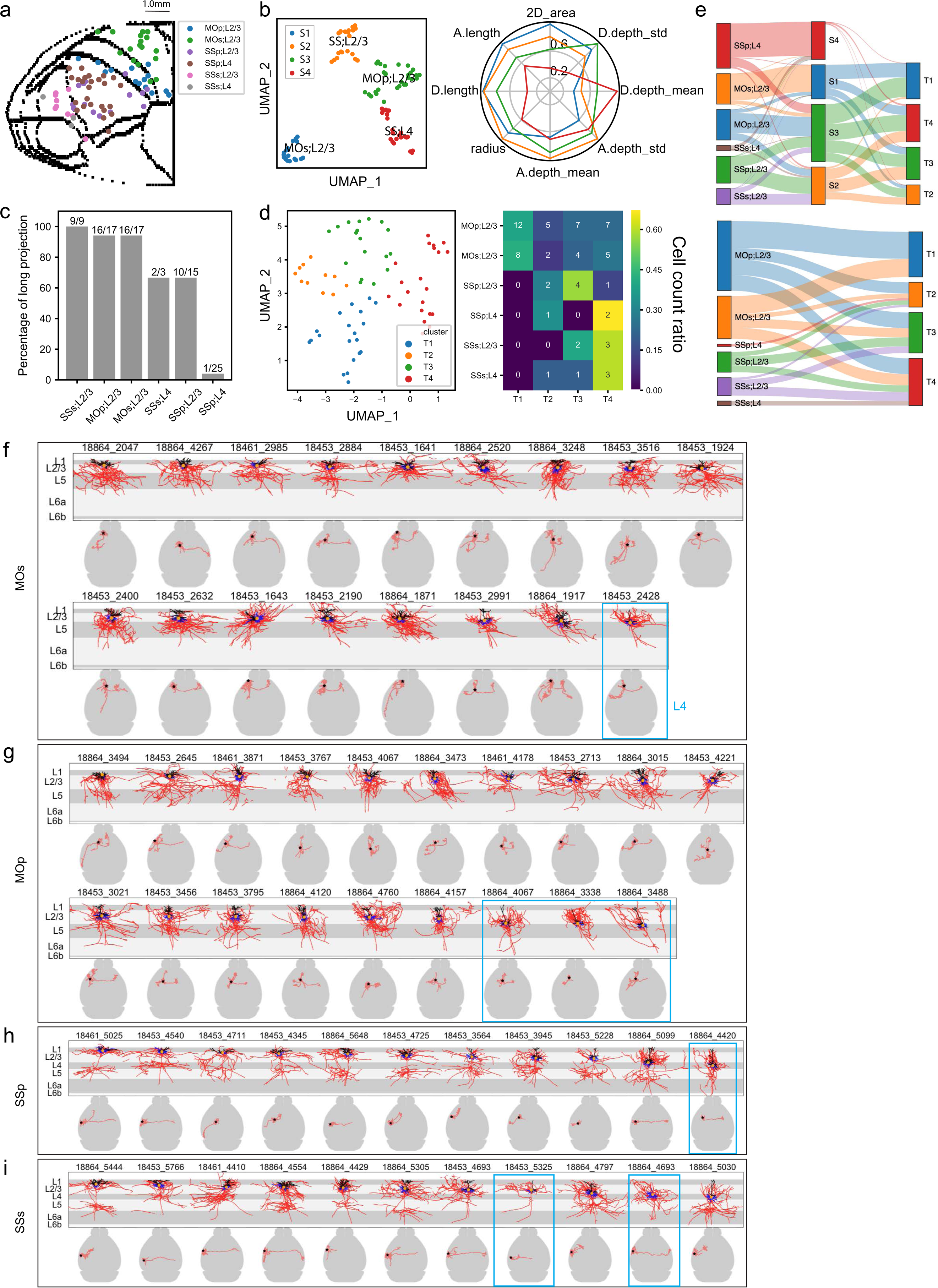
Local morphological and long-range projection analysis of cortical L2/3/4 IT neurons. **a**, Cortical surface flatmap showing the soma locations of reconstructed L2/3/4 IT neurons from MOp, MOs, SSp and SSs. **b**, Clustering based on local dendritic, axonal and soma location features divides L2/3/4 IT cells into 4 clusters. UMAP dimension reduction was performed, followed by k-means clustering using UMAP embeddings as input features. Clustering results shown by the UMAP representation and polar plot of main features. **c**, Percentage of cells from each region and each layer that have long-range projections. **d**, Clustering based on long-range projection targets, shown by the UMAP representation and confusion matrix of soma location and projection clusters. Clustering is performed by UMAP embedding and k-means. **e**, Sankey plots showing the correspondence among soma locations, local clusters and long-range projection clusters for individual neurons. **f-i**, Comparison of local morphologies (upper panels; apical dendrite in black, basal dendrite in blue, axon in red, soma as an orange dot) and whole-brain projections (lower panels; axon in red, soma as a star) for MOs (f), MOp (g), SSp (h) and SSs (i) neurons. Neurons are ordered based on the depths from pial surface of their somas. L4 IT neurons are marked by the blue boxes. The others are L2/3 IT neurons.

**Figure 6.**
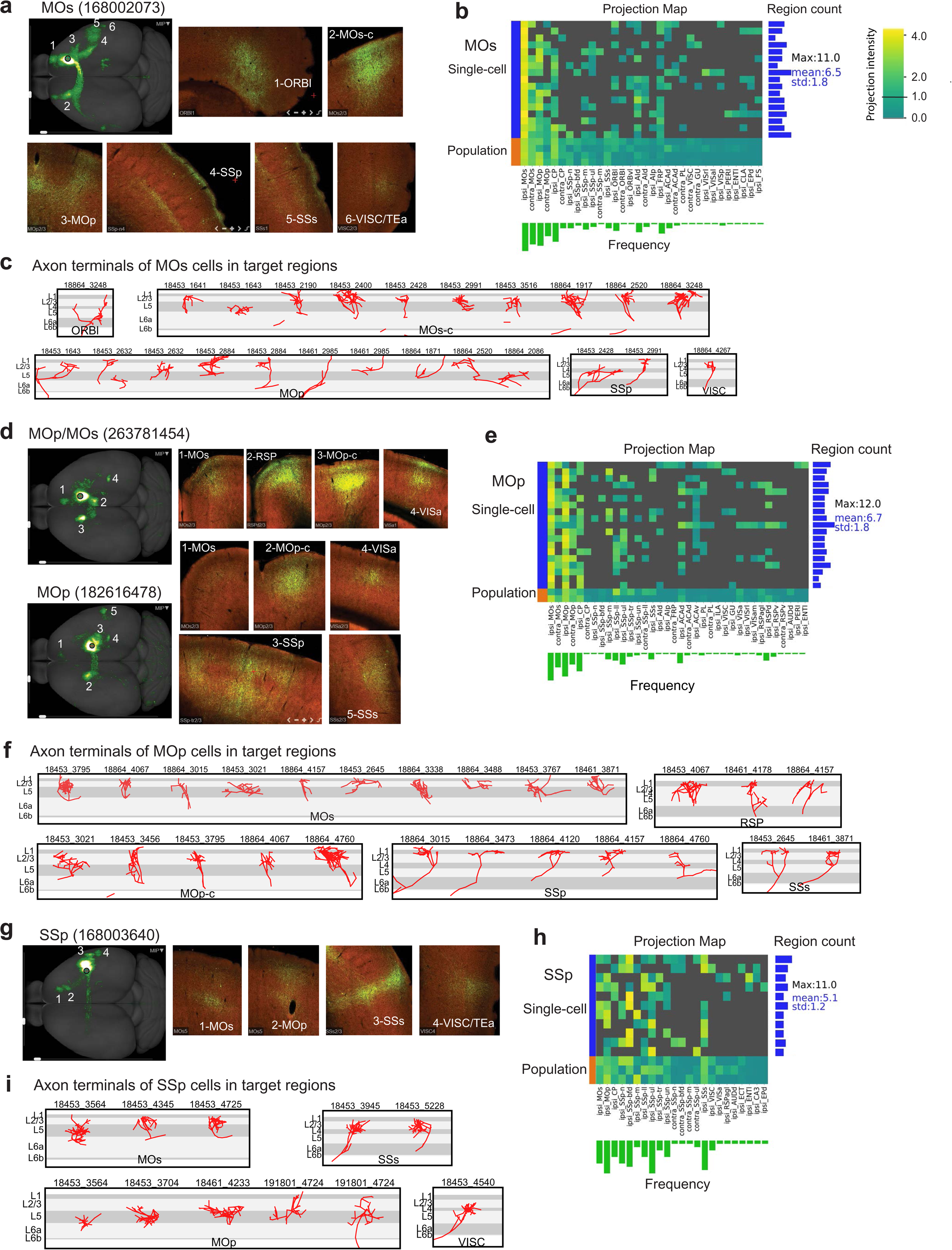
Comparison of long-range projection patterns between individual cortical L2/3/4 IT neurons and mesoscale population-level projections. **a**, A representative mesoscale experiment from MOs, shown in a maximum projection whole-brain top-down view and individual higher power images showing axon termination patterns in major target regions. **b**, Projection matrix of MOs single cells and population-level mesoscale experiments (168002073, 266645328, 272822110 and 587659400) along with some basic statistics for the single cells (i.e. target region counts per cell, frequencies per target region). Projection strengths are scaled to 0-4 and visualized with a cutoff at 1 to minimize spurious false positives due to issues such as imaging artifact (in mesoscale experiments), imprecise registration and minor passing fibers. Cells are ordered by soma depth from pial surface. Only brain regions targeted (relative projection strength >1) by any single/mesoscale samples are included. **c**, Axon terminals of MOs neurons in specified target regions. Because not all neurons project to all target regions, all detected axon terminals from any neurons for each target region are shown here. MOs-c, contralateral MOs. **d-f**, Same panels as a-c but for MOp. Two representative mesoscale experiments (182616478 and 263781454) from MOp are shown in d, which are also used as population projections in e. **g-i**, Same panels as a-c but for SSp. Mesoscale experiments used as population projections in h are 168003640, 298830161 and 278317945.

As different thalamic regions have unique cortical projection patterns ^16, 57, 59^, we first compared projections of individual thalamic neurons with projections of anterograde bulk tracing from the Allen Mouse Connectivity Atlas (**Supplementary Table 3**). We find that single cell projections are basically consistent with the bulk anterograde tracing results for the thalamic nuclei their somas reside in (Fig. 4a-h, left two columns).

Also consistent with prior knowledge, we find that the single sensory-motor relay neurons usually have one major axon arbor targeting the primary sensory or motor cortex of the corresponding modality, i.e. VPM and VPL to primary somatosensory area (SSp), VPMpc to gustatory areas (GU), VPLpc to visceral area (VISC), LGd to primary visual area (VISp), MG to auditory areas (AUD), and VAL to primary motor area (MOp). Axons from these nuclei terminate predominantly in L4 and L6, consistent with the “core-type” classification (Fig. 4a,b,d, Extended Data Fig. 5). Reconstructed neurons from AM, SMT and posterior MD also send a single major axon arbor to various parts of orbital area (ORB), with a similar mid-layer termination pattern, suggesting these neurons also belong to the “core” projection type (see also Harris et al. 2019) (Fig. 4c,e, Extended Data Fig. 5).

We quantitatively analyzed morphometric features of 1,103 axon arbors from 624 neurons located in the primary sensory thalamic nuclei: VPM, VPL, LGd and MG (Extended Data Fig. 7). Our results suggest there are two major types of axon arbors targeting cortical layer 4 (spanning L4 and lower L2/3); a smaller “type 1” arbor and a larger “type 2” arbor (cortical area >0.3 mm^2^). We also identified a “type 3” arbor terminating in cortical L6; this type was most often a minor branch originating from the type 1 or type 2 arbors, so we did not use it to classify neurons. Single neurons were thus assigned to either small-arbor or large-arbor type (Fig. 4a,b,d). The ratio of large- to small-arbor neurons varies by thalamic region. Large-arbor neurons account for 28.3% of the total reconstructions from VPM, 28.0% from VPL, 46.8% from LGd and 58.3% from MG. Notably, somas with small and large arbors are spatially intermingled in each nucleus (Extended Data Fig. 7e). Neurons located in SMT are also separable into small- and large-arbor types like the primary relay nuclei (Fig. 4c), whereas the current set of AM neurons all have small arbors and the posterior MD neurons all have large arbors (Fig. 4e, Extended Data Fig. 5).

**Figure 7.**
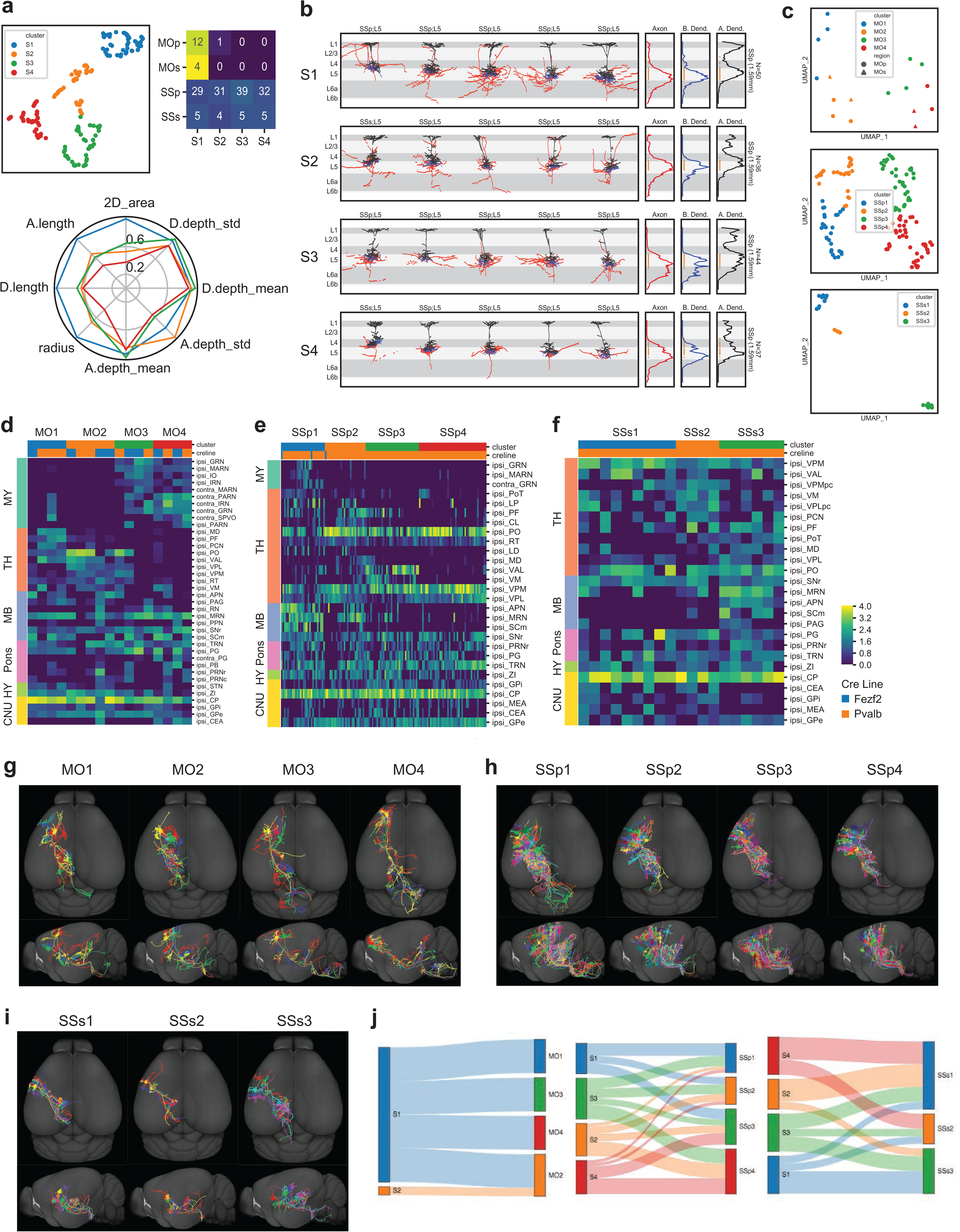
Local morphological and long-range projection analysis of cortical L5 ET neurons. **a**, Clustering based on local dendritic, axonal and soma location features divides L5 ET cells into 4 clusters, as shown by the UMAP representation, soma-cluster confusion matrix and polar plot of main features. Clustering approach is the same as used for Fig. 5b. **b**, Local morphologies of example neurons and average vertical profiles for each local cluster. Broken lines are due to the substantially tilted nature of some of these neurons. **c**, Clustering based on long-range projection targets, carried out separately for each region (MOp/MOs combined, SSp and SSs), as shown by the UMAP representations. **d-f**, Projection matrix heatmaps for MOp/MOs (d), SSp (e) and SSs (f) neurons, showing representative target brain regions of each neuron. Columns represent single cells sorted by cluster assignments. Rows represent target regions. **g-i**, Whole-brain projection overview of individual neurons in each cluster for MOp/MOs (g), SSp (h) and SSs (i). **j**, Sankey plots showing the correspondence between local clusters and long-range projection clusters for individual neurons.

A small fraction of the core-type thalamic projection neurons have more than one axon arbor targeting different cortical areas (6.77% for VPM, 14.44% for VPL, 12.5% for MG, but 0% for LGd). In the case of these cells in VPM and VPL, usually they have a larger main arbor targeting SSp, and a smaller secondary arbor targeting the supplemental somatosensory area (SSs) (Fig. 4a,b). MG neurons with two or more cortical targets are mostly of the large-arbor type (Fig. 4d). These multi-target MG neurons are more like the matrix type thalamocortical neurons described below, showing stronger projections to L1 and L5, located in the associational parts of MG (*e.g.*, MGm) medial to the core relay auditory nucleus, MGv.

Outside the sensory-motor relay thalamic nuclei, nearly all reconstructed neurons have a large diffusely branched axon arbor and/or several arbors projecting to multiple cortical areas, often with columnar or L5-dominant axon termination patterns (Fig. 4e-h, Extended Data Fig. 6). Many (81%) of these cells also have axon branches >1 mm long in L1, consistent with the “matrix” type, but they also exhibit a diverse range of projection and morphological patterns. For example, LP neurons preferentially project to two or more higher visual cortical areas. They do not directly project into VISp, only one out of 16 reconstructed LP neurons has axon fibers in L1 extending into VISp. LP neurons can be roughly divided into an anterior and a posterior group, consistent with previous functional studies ^60^. Posterior LP neurons mainly project to lateral and posterior higher visual areas, whereas anterior LP neurons mainly project to medial and anterior higher visual areas with some extending an axon projection into anterior cingulate area (ACA) (Fig. 4f, Extended Data Fig. 6). PO neurons project to both SSp and MOp/MOs (secondary motor area). Their axon arbors in these target regions terminate broadly across layers with an apparent preference in lower L2/3, with 5 out of 7 sending rich axon arbors (>1 mm) to L1 (Fig. 4g, Extended Data Fig. 6). Neurons in anterior MD appear very different from those in posterior MD and are more similar to those in neighboring nuclei such as IAD and CM. They have multiple axon arbors that target multiple medial and lateral prefrontal cortical areas including prelimbic area (PL), ORB and agranular insular area (AI) (Fig. 4e, Extended Data Fig. 6). VM neurons have multiple axon arbors, heavily targeting MOp/MOs with additional branches targeting various somatosensory areas (Fig. 4h, Extended Data Fig. 6).

Quantitative interareal projection matrix (Extended Data Fig. 8, **Supplementary Tables 2 and 3**) confirms the above observations and further demonstrates the distinction between core- and matrix-type neurons, with the former (from VPM, VPL, VPMpc, VPLpc, LGd, VAL, AM, and SMT) predominantly targeting a single cortical area (sometimes with a secondary area) and the latter (MG, MD, LP, PO, LD, VM, RE, and CM) targeting multiple cortical areas. Note that MG contains a mixture of core and matrix cells. Within each nucleus, on the other hand, individual neurons show a high degree of consistency with each other in projection patterns, and each neuron’s projection pattern is also similar to the population projection pattern for that nucleus.

**Figure 8.**
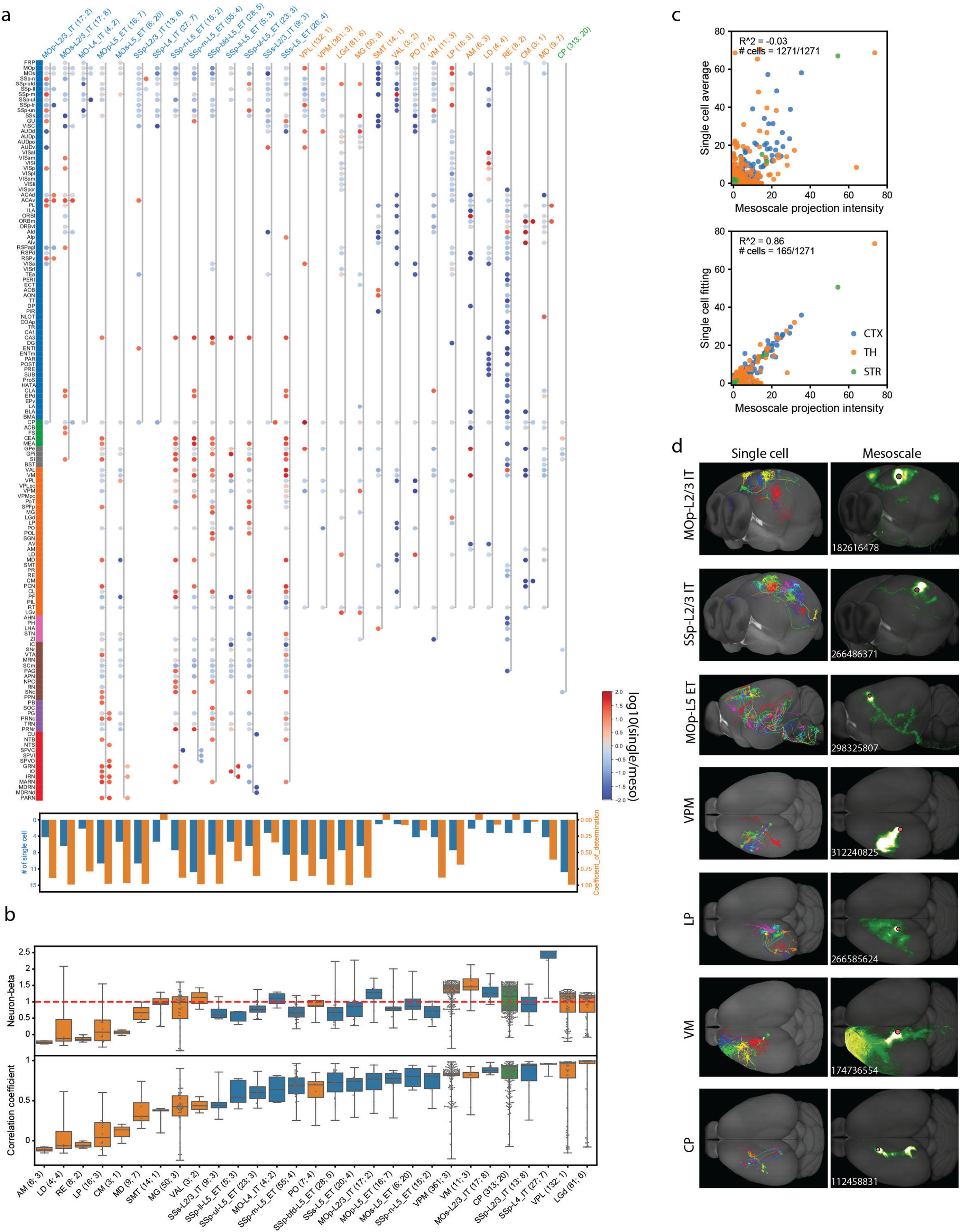
Combination of single neuron morphologies recapitulates population-level mesoscale projection patterns. **a**, Comparative projection map of single cell and mesoscale data. Individual samples are grouped by brain areas and/or cortical layers based on soma locations (single cell) and injection sites (mesoscale). Each group is represented by a stretch of connected dots with ipsilateral and contralateral targets on the left and right hemisphere, respectively. Projection intensities are quantified as log(percentage+1). Selected regions are defined at a cutoff 0.5 and targeted by at least 50% mesoscale or 10% single cells. Dot colors are scaled by the log10 of single cell and mesoscale strength ratio. (lower panel) Coefficients of determination (orange bars) and number of cells (blue bars) of mesoscale regression by single cells (described in c). **b**, Boxplots of neuron-beta and correlation coefficients between single cells and group-average of mesoscale data. Individual comparisons shown as swarm plots overlapped with boxes. The first and second numbers in the group labels in a and b indicate the numbers of single cells and mesoscale experiments, respectively. **c**, Approximation of mesoscale projections by single cell projection strengths (1,271 cells used) by group-average (upper) or by linear regression (lower). LASSO regularity was applied during regression, to reduce the number of single cells with non-zero weights (representative cells). **d**, Visualization of projection patterns constituted by representative cells and mesoscale projection intensities.

### Cortical L2/3/4 intratelencephalic (IT) neurons have variable intracortical projection targets

We reconstructed 160 cortical neurons in total from the Cux2-CreERT2 line, and further analyzed 93 of these neurons in somatosensory and motor regions: SSp (n=40), SSs (n=15), MOp (n=20) and MOs (n=18) (Fig. 5a, **Supplementary Table 2**). *Cux2*+ neurons are located in L2/3 or L4 and their long-range projections are confined within cortex and striatum, consistent with them belonging to the corticocortical projecting IT subclass ^61^. Clustering analysis using local dendritic and axonal morphological features grouped these cells largely by region and layer (Fig. 5b).

We identified 25 cells from SSp and 3 cells from SSs to be in L4. These cells were distinguished from L2/3 neurons using morphological features; they have either no apical dendrites (*i.e.*, spiny stellate cells) or a simple apical dendrite that does not branch in L1 (*i.e.*, untufted or star pyramid cells), in contrast to the pyramidal L2/3 cells which have tufted apical dendrites in L1 ^61^ (Fig. 5h-i, Extended Data Fig. 9). L2/3 cells also have local axons in L2/3, which also project downward into L5, whereas L4 cells have local axons mainly projecting up to L2/3, another differentiating feature ^62^. Interestingly, we also found 4 neurons from MOp/MOs with these L4-like features – minimal apical dendrites and upward-projecting local axons (Fig. 5f-g), and they are located between L2/3 and L5 since L4 is not delineated in CCFv3, suggesting that these are the L4-like cells located in motor cortex ^63^ that can also be identified transcriptomically (see flagship paper).

**Figure 9.**
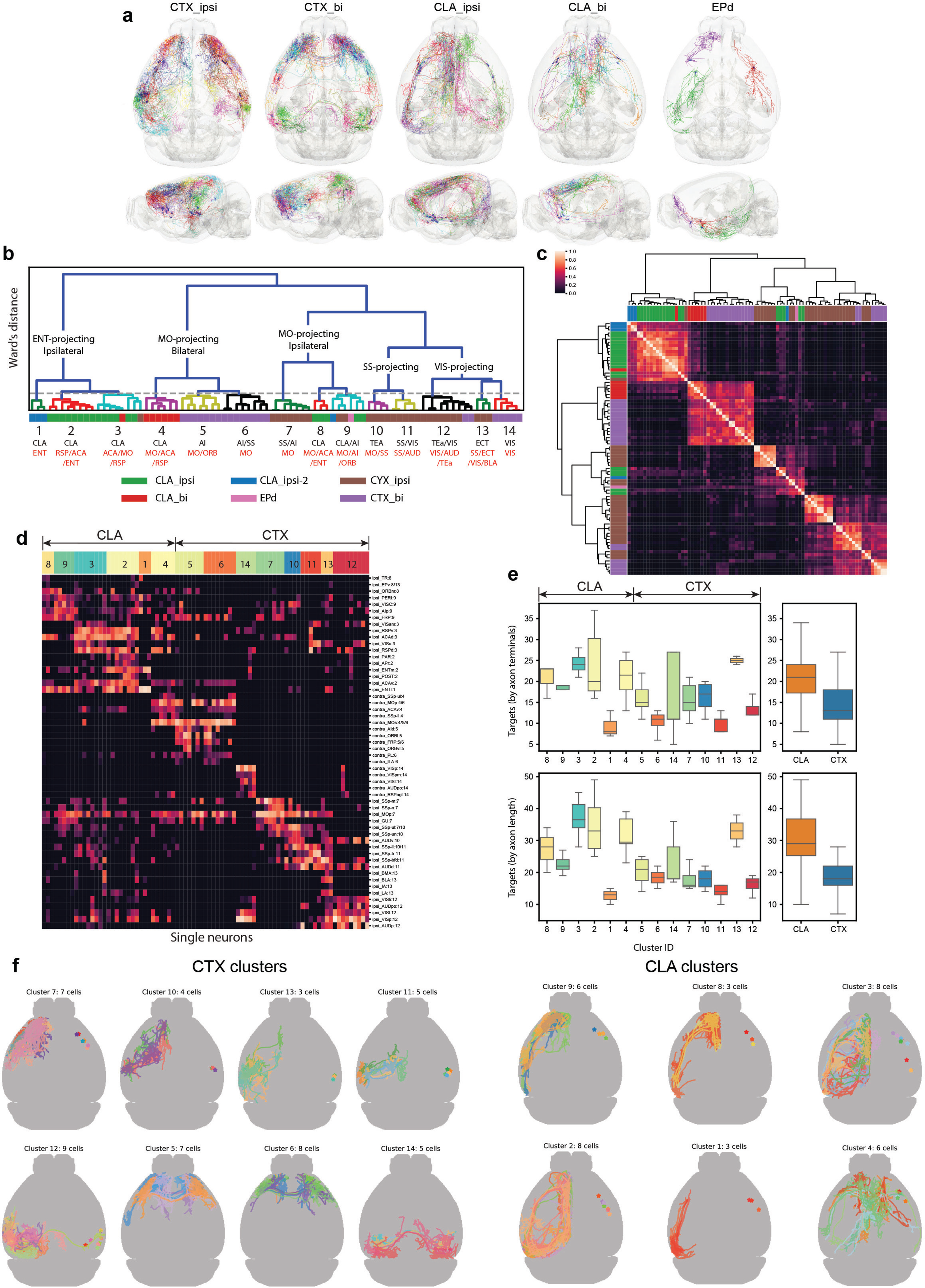
Extensive projection diversity of the L6 Car3 subclass of cortical and claustral neurons. **a**, Individual L6 Car3 neurons are shown in manually divided groups: CTX_ipsi (n=33), cortical neurons projecting to ipsilateral cortex only; CTX_bi (n=26), cortical neurons projecting bilaterally; CLA_ipsi (n=24), claustral neurons projecting to ipsilateral cortex only; CLA_bi (n=10), claustral neurons projecting bilaterally; EPd neurons (n=3) form a distinct morphological group, with specific projections to olfactory areas and limbic cortical areas. **b-c**, Integrated co-clustering dendrogram (b) and matrix (c) obtained by averaging the co-clustering matrices of four feature sets: projection pattern, soma location, axon morphology and dendrite morphology. Threshold for cluster calls is shown as the dashed line. Each cluster is annotated by the brain regions where somata (black) and axon clusters (red) reside. Regions were selected to represent >50% of cluster members. Side bars indicate manually assigned types with color codes shown below the matrix. **d,** Projection matrix heatmap for representative target brain regions of each neuron. Columns represent single cells sorted by cluster assignments. Rows represent targets, and the number following each target name indicates the dominant cluster ID for the row. **e,** Total number of cortical targets innervated by each neuron grouped by clusters. Ipsilateral and contralateral targets are counted separately. A minimum of one axon terminal (top panel) or 1,000 µm of axon length (bottom panel) is used as the threshold to label a region as “targeted”. **f,** Top-down views of neurons in each cluster. Neurons are all flipped to the left hemisphere for comparison of axon projection patterns. Stars indicate soma locations and are flipped to the right hemisphere for visualization purpose.

Several regional differences in axon projections of these L2/3/4 IT neurons are observed. The proportion of cells with intracortical long-range projections (defined as having >5 mm axon branches that are >1 mm away from the soma) is larger in SSs, MOp and MOs than SSp (Fig. 5c). Consistent with prior notion ^61^, all but one SSp L4 cells have only local axons but no long-range projections (Extended Data Fig. 9). However, nearly all L4 and L4-like cells in SSs, MOp and MOs do have axon projections outside of their local area (Fig. 5f-i), as we reported before ^57^. For the cells with long-range projections, the average total length of long-projecting axons and the number of target regions per cell are also greater in MOp/MOs than in SSp/SSs (p < 0.003 for axon length and <0.002 for target regions, Mann-Whitney rank test).

Several recent studies have shown that transcriptomically defined cortical IT neurons are organized by layer, but also exhibit a gradual transition in transcriptomic type spatially along the cortical depth, by scRNA-seq ^64^, MERFISH ^65^ and Patch-seq ^66^. Here we arrange the L2/3/4 IT cells according to the depth of their soma from the pial surface, region by region, and examine their local dendritic and axonal morphologies and long-range projections (Fig. 5f-i). We find that within each region, across depths individual neurons exhibit variable long-range projection patterns with no obvious depth correspondence, suggesting that long-range projection patterns of these L2/3/4 IT neurons may not correlate with their transcriptomically defined subtypes.

Clustering based on long-range projection patterns grouped the cells into 4 clusters (T1-T4), although variation appears largely continuous (Fig. 5d). We did not observe a clear one-to-one correspondence between these long-projecting clusters and either local morphology clusters (S1-S4) or soma areal origin (Fig. 5e). Comparing single neuron and population projection patterns (**Supplementary Table 3**) shows that all neurons together recapitulate the population projection pattern, but each neuron selects a subset of projection targets (Fig. 6a-b, d-e, g-h), in contrast with the thalamic neurons reported above. Thus, the selection of a subset of projection targets by each neuron appears random without specific correlation to soma depths or dendritic morphologies.

We next asked if axon termination patterns of single cells recapitulate the overall feedforward or feedback projection patterns apparent at the population level ^57^. Given that each L2/3/4 IT cell projects to only a subset of their intracortical targets, we pooled all the terminal axon arbors to a specific target from all the cells within a source region (Fig. 6c,f,i). Interestingly, we find that nearly all axon terminals of SSp cells are concentrated in middle layers (L2/3-5) of all target regions (MOs, MOp, SSs and VISC, Fig. 6i), suggesting all outward projections from SSp are feedforward, consistent with it being at the bottom of the cortical hierarchy ^57^. On the other hand, axons of MOp cells mainly terminate in L2/3-5 of target areas MOs and contralateral MOp, but have prominent termination in L1 of target areas SSp and SSs (Fig. 6f), consistent with the notion that the former projections are feedforward and the latter feedback. The MOs cells do not show such a clear division with the available number of terminal axon arbors to different targets (Fig. 6c).

Finally, we investigated the projection target specificity of transcriptomic cell types using Retro-seq ^4^, in which the transcriptomes of 822 retrogradely labeled neurons from SSp, SSs, MOp and MOs were mapped to our transcriptomic taxonomy ^64^ to identify the transcriptomic type of each retrogradely labeled neuron (Extended Data Fig. 10, **Supplementary Table 4**). We found that for each source region, L2/3 and L4/5 IT neurons labeled from different injection targets were mostly mapped to a few common transcriptomic types. Taken together, the above results suggest that within the L2/3/4 IT subclasses, the projection patterns at a single cell level do not correlate one-to-one with the cell’s transcriptomic type in the adult animal.

### Cortical L5 extratelencephalic (ET) neurons show distinct subcortical projection specificity

We reconstructed 251 L5 ET neurons in total from two Cre lines: Fezf2 and Pvalb, and analyzed more than half of these neurons from the SSp (n=130), SSs (n=19), MOp (n=13) and MOs (n=4) (Extended Data Fig. 11, **Supplementary Table 2**). We first clustered the cells into four groups based on their local dendritic and axonal features (Fig. 7a-b). Cluster S1 is the most unique; these neurons have larger dendritic and local axonal arbors and nearly all MOp and MOs neurons belong to this cluster. Clusters S2-S4 are separable by features that capture the complexity of local axons (*e.g.,* length, total area containing axon, and axon lamination).

The L5 ET neurons have 11.3 projection targets (defined using a stringent threshold of axon length >1 mm in the target region) on average, significantly more than the L2/3/4 IT neurons which have 8.0 projection targets on average (p<2.0e-6, Mann-Whitney rank test). Like L2/3/4 IT neurons, L5 ET neurons exhibit extensive diversity and heterogeneity in each of their selected subset of projection targets. Nonetheless, we clustered these cells based on their long-range projections within each region (MOp/MOs combined, SSp, and SSs) and identified subtypes with specific projection targets (Fig. 7c-f).

L5 ET neurons in MOp/MOs form 4 distinct projection types (Fig. 7d,g). Notably, MO3 and MO4 types project to regions in the medulla whereas MO1 and MO2 types do not, a distinction consistent with recent findings from combined transcriptomic/epigenomic and projection studies ^32, 67^. Note that few SSp neurons and no SSs neurons project to the medulla (Fig. 7e,f), suggesting medulla projection may be primarily a feature of MOp/MOs neurons. Within the medulla-projecting types, MO4 neurons have few projections to other parts of the brain such as thalamus and basal ganglia compared to MO3 neurons. MO4 neurons also have stronger preference to project to the contralateral side of medulla. Within the non-medulla-projecting types, MO1 and MO2 neurons again display differential projection patterns in the thalamus, with MO1 preferentially targeting VM/VAL/MD whereas MO2 preferentially target PO.

L5 ET neurons in SSp can also be divided into 4 clusters based on their projections (Fig. 7e,h). The 4 clusters are most distinguishable by their differential targeting of various thalamic nuclei. For example, SSp3 neurons have more projections to VAL and VM than neurons in other clusters. In addition, cluster SSp1 contains most of the cells projecting to medulla. Cells in cluster SSp4 have the fewest subcortical projections. In SSs, L5 ET neurons can be divided into 3 clusters (Fig. 7f,i). Cells in clusters SSs1 and SSs2 have fewer projections to the midbrain compared to those in SSs3. On the other hand, SSs1 and SSs2 cells project to CEA and MEA but SSs3 cells do not.

Similar to the IT neurons, there is no clear one-to-one correspondence between local morphology and long-range projections of L5 ET neurons, except that neurons in the medulla-projecting SSp1 cluster are mostly present in the S1 and S3 clusters, which have deeper soma depth and larger cortical areas (Fig. 7j). This is consistent with previous finding that these neurons are located in the deeper part of layer 5 ^32^. We compared the local morphologies of MOp/MOs neurons with or without medulla projection and found that medulla-projecting neurons have a weak tendency for more extensive and complex dendrites (Extended Data Fig. 12).

### Combination of single neuron projections recapitulates population-level mesoscale projection pattern

Mapping axonal projections at the mesoscale has revealed region-specific patterns of connectivity across the mouse brain ^14, 16, 57^, yet how projection motifs at the single cell level compose the population-level patterns are still unclear for most of the brain regions. To directly compare the mesoscale and single cell projection patterns, we identified 1,271 single cell morphologies and 141 mesoscale experiments from the Allen Mouse Brain Connectivity Atlas, matched based on soma or injection site being within the same CCFv3 structure (**Supplementary Table 3**). This dataset covers 14 cortical areas and layers combined, 14 thalamic nuclei and one striatal structure (CP). Using this location-matched dataset, we determined the targets of single cells and mesoscale groups by thresholding at both individual and group levels and constructed a comparative map (Fig. 8a, see Methods). Overall, the combined single cell projection pattern from a region (and cortical layer) is highly concordant with that of the mesoscale experiments.

We also observe a few exceptions to this general trend. The combined patterns from single L5 ET neurons across several cortical areas collectively project to more subcortical targets than mesoscale experiments (indicated with more red colored dots), likely due to broader distribution of the single neurons in the source areas. On the other hand, for several thalamic nuclei (i.e., SMT, VAL, AM, LD, RE and CM), single neurons collectively have not captured the full projection patterns from mesoscale experiments (indicated with more blue dots). This difference could be due to several reasons: (1) since some of these nuclei are small, the mesoscale experiments may include projections labeled from neighboring nuclei so the single cell data may more accurately represent the true output pattern; (2) the number of reconstructed single neurons is still relatively small and may not fully represent all projection types in a given nucleus; (3) the reconstructed neurons may represent only a subset of the cell types located in these nuclei given the Cre driver labeling method employed, and there may be other types of projection neurons not labeled in these specific Cre lines.

To quantitatively compare the single cell and mesoscale tracer experiments, we calculated the correlation coefficient of each single cell’s brain-wide projection weights with the average projection weights from the location-matched mesoscale experiments (Fig. 8b). The correlation coefficient ranges from −0.11 (e.g. AM) to 0.99 (e.g. LGd), with a median of 0.82. High correlation coefficients may indicate simple compositions of projecting patterns, e.g. LGd with almost pure VISp projections and CP projecting to either GPe or SNr. Low correlation coefficients may indicate complex composition of projecting patterns, e.g. CM, AM and RE for reasons mentioned above. To compare single cell projection strength relative to mesoscale data, we developed a ‘Neuron-beta’ metric, as the covariance of a single cell and the average of mesoscale samples, relative to the mesoscale variance (Fig 8b, see Methods). Single cells with Neuron-beta values >1.5 correlate well with mesoscale data but fluctuate more variably. For example, individual VM neurons are highly diverse but positively correlated with mesoscale data. Small (<0.5) positive Neuron-beta values result from low correlation (<0.13) of single and mesoscale data. For cell types with Neuron-beta values around 1, single cell and mesoscale data appear to be comparable.

To study how well the mesoscale projection pattern could be broken down to our set of single cells, we performed linear regression with Lasso regularization. This approach selects a minimal set of single cells and uses weighted summation of single cell axon length to approximate the cell type specific mesoscale axonal weights. The overall coefficient of determination (R^2^) is 0.86, indicating that mesoscale connectivity is recapitulated well (Fig. 8c). R^2^ is >0.8 for most groups except for the above-mentioned thalamic nuclei and L4 IT neurons from SSp (Fig. 8a lower panel). The discrepancy in the SSp L4 IT group is likely due to the fact that the Cre lines of L4 mesoscale experiments (Nr5a1-Cre, Scnn1a-Tg3-Cre and Rorb-IRES2-Cre) are not entirely L4 specific and also label some L5 cells ^4^, thus the single cell projection pattern is more accurate in this case. Only 165 out of 1271 single cells with non-zero values contribute to the regression. These cells represent a minimal set of stereotypes to make up the population level connectivity (Fig. 8d). Averaging across all single cells shows a low level of approximation (R^2^=-0.03), suggesting highly diverse morphologies and projection patterns among the single cells.

### Extensive projection diversity in transcriptomically homogeneous Car3 cortical and claustral neurons

Finally, we investigated a special type of cortical excitatory neurons, the L6 Car3 IT transcriptomic subclass ^4, 64^, whose morphology and projection patterns have been unknown. This subclass of neurons is selectively labeled by a unique set of marker genes including *Gnb4*, which is located in the deep layer (mostly L6) of all lateral cortical areas and shares the same transcriptomic clusters with neurons from the claustrum (CLA, Extended Data Fig. 13a,b). Previous mesoscale anterograde tracing showed that claustrum neurons project widely into cortex, with particularly strong connections with prefrontal and retrohippocampal cortical areas ^68^. We performed similar experiments with the same AAV tracer injected in CLA, SSs or SSp of the Gnb4-IRES2-CreERT2 mice, and found that L6 Car3 neurons in SSs and SSp also showed intracortical projections, but with a more restricted, distinct set of targets compared to CLA projections (Extended Data Fig. 14).

We reconstructed 96 neurons from the cortex and CLA of the Gnb4-IRES2-CreERT2 line (**Supplementary Table 1**), which contain 34 CLA neurons, 59 neurons from multiple lateral cortical areas, and 3 neurons from the dorsal part of endopiriform nucleus (EPd) (Fig. 9a, **Supplementary Table 2**). We performed clustering using four feature sets – projection pattern, soma location, axon morphology and dendrite morphology, and identified 14 clusters (Fig. 9b-d). We calculated the total number of projection targets using two different thresholds to label a region as “targeted” (Fig. 9e). Using a minimum of 1 mm of axon length (identical threshold as used in ^22^) we found the median number of targets to be 29 for CLA neurons and 18 for cortical L6 Car3 neurons. Alternatively, using the existence of at least one axon terminal as the threshold we found the median number of targets to be 21 for CLA neurons and 12 for L6 Car3 neurons. In both cases, the numbers of targets were substantially greater than the above L2/3/4 IT neurons as well as that reported previously for L2/3 IT neurons of the primary visual cortex ^22^.

All CLA and cortical L6 Car3 neurons project almost exclusively into the cortex with none or minimal axon projections into the striatum. This is another major difference between the L6 Car3 cortical and claustral neurons and other types of corticocortical-projecting IT neurons which have substantial axon collaterals projecting to the striatum ^61^. EPd neurons have distinct projections to frontal cortex, piriform cortex and subcortical olfactory areas ^69, 70^.

Cortical L6 Car3 cells are assigned to 8 clusters, with clusters 7, 10, 11, 12 and 13 projecting ipsilaterally, and clusters 5, 6 and 14 bilaterally (Fig. 9f). The clusters are arranged topographically from anterior to posterior cortex based on both soma location and projection target specificity; each cluster contains a group of neurons that are located close to each other and project to similar cortical target areas. Clusters 1, 2, 3 and 8 belong to the CLA-ipsilateral group, in which cluster 1 is lateral-projecting and clusters 2, 3 and 8 are midline-projecting (Fig. 9f). Cluster 4 contains CLA-bilateral midline-projecting cells. Cluster 9 contains a mixture of CLA, EPd and cortical cells, as they show similar soma locations and axon projection features with preferential projection to the frontal pole (FRP) and ORBl areas on the ipsilateral side.

Interestingly, we identify a unique group of four L6 Car3 cells, all with their somas located in temporal association areas (TEa) and ectorhinal area (ECT), three of which belonging to cluster 13 whereas the 4^th^ one is an outlier, have substantial projections into amygdala areas including lateral amygdalar nucleus (LA), basolateral amygdalar nucleus (BLA) and basomedial amygdalar nucleus (BMA), in addition to their cortical targets (Fig. 9b,d). There are also several CLA neurons, scattered in several clusters, with minor axon collaterals projecting into amygdala areas ^21, 68^ (Fig. 9d).

In our single-cell transcriptomic taxonomy of the entire mouse cortex and hippocampus ^64^, the L6 Car3 subclass contains 1,997 cells, of which 799 are from CLA and 1,198 are from various cortical regions (Extended Data Fig. 13a,b, **Supplementary Table 4**). Cells from different regions are distributed similarly across 5 clusters within the subclass, suggesting a lack of region-specificity for any cluster and that these cortical and claustral cells are highly related to each other, possibly reflecting common or closely related developmental origins. In an attempt to link molecular identities with the projection diversity described above, we performed Retro-seq on cells isolated from CLA (238) and cortical areas SSs (11) and TEa (35) that were labeled by retrograde tracers injected into far apart cortical areas, ACA (medial), MOp (central), ORBl or VISpl (lateral) (Extended Data Fig. 13c,d, **Supplementary Table 4**). The CLA and cortical Retro-seq cells projecting to different cortical areas share the same set of L6 Car3 clusters, indicating no clear one-to-one correspondence between transcriptomic clusters and projection target specificity.

Taken together, the remarkable morphological diversity of these fully reconstructed neurons from claustrum, endopiriform nucleus and cortex, which all belong to a single, transcriptomically defined L6 Car3 subclass, demonstrates that long-range axonal projections vary greatly according to their cell body locations, indicative of a combined functional and topographic organization of structural connectivity.

## DISCUSSION

To fully understand the morphological and projection diversity and specificity of neurons across the brain, a large number, likely in the range of hundreds of thousands of neurons will need to be examined. Approaches such as MAPseq ^22, 71, 72^ can quickly survey projection specificity at the regional level for many neurons in a high throughput manner. However, many essential details can only be obtained through full morphological reconstructions. Collecting such ground truth data provides an invaluable opportunity to uncover principles of neuronal diversity and circuit organization, informing functional studies. To this end, we here report a large set of full neuronal morphologies, generated from a standardized platform we established, which reveal region- and cell type-specific diversities.

Our unique labeling strategy using stable and universal transgenic reporter mouse lines coupled with a variety of sparse Cre delivery methods has several advantages. First, the TIGRE2.0-based transgenic reporter lines, especially Ai166 which expresses a farnesylated GFP, produce very bright GFP labeling of axon fibers under fMOST imaging, revealing numerous terminal boutons, an essential requirement for obtaining truly complete morphologies. Second, this strategy enables sparse labeling across multiple regions within the same brain, improving efficiency compared to other methods (*e.g.*, *in vivo* electroporation or stereotaxic virus injection). Third, the labeling is highly consistent from cell to cell, cell type to cell type, region to region, and brain to brain, reducing variability and enhancing reproducibility. Finally, sparse Cre recombination can be achieved through the use of transgenic Cre or CreERT2 driver lines labeling any neuronal type, or low-dose Cre viral vectors delivered through either local or systemic (*e.g.*, retroorbital) injections.

Development of novel and accessible software tools are essential for reconstruction efficiency. Such powerful tools have been recently developed by the Janelia MouseLight project ^36^. Similarly, our enhanced Vaa3D-based reconstruction toolkit streamlines large-volume fMOST image data processing and computation-assisted manual reconstruction. The registration of the fMOST whole-brain datasets to the CCF allows quantification of projection strength in each target region across the entire brain for each neuron, and subsequent data-driven clustering to identify similarities and differences across neurons and to group them into types. The accumulation of an increasingly larger set of fully reconstructed neurons in the future can be used as training datasets to develop machine learning-based automatic reconstruction algorithms to further boost the throughput of reconstruction.

Our extensive and detailed analysis of this large collection of reconstructed neurons has yielded a number of novel findings regarding neuronal projection diversity at multiple organizational levels. First, at the regional level, neurons from different brain regions follow different convergence or divergence rules in their long-range projections. Striatal neurons, both GPe-projecting and SNr-projecting types, have highly convergent projections into their respective targets. The core-type thalamic neurons have largely point-to-point projections to their cortical targets. The matrix-type thalamic neurons, the claustral neurons and all classes of cortical neurons have divergent projections to a few (for thalamic neurons) or many (for claustral and cortical neurons) target regions. The degree of similarity between individual neurons within a given region also varies across regions. The striatal neurons and core-type thalamic neurons have one dominant axon branches that are highly consistent among individual neurons within the same type. The matrix-type thalamic neurons usually have a few axon branches but are also mostly consistent among the individual neurons within each nucleus. The cortical and claustral neurons, on the other hand, have highly variable axon branching and projection patterns with each neuron selecting only a subset of the targets.

Second, at the major cell class or type level, robust distinctions in morphological features between types have been discovered. For example, striatal GPe-projecting neurons have more extensive local axon arborization than SNr-projecting neurons. The core-type thalamic neurons can be divided into small-arbor and large-arbor types, whereas the matrix-type thalamic neurons all have much larger, diffusely distributed axon arbors. In the cortex, the L2/3/4 IT, L5 ET and L6 Car3 subclasses exhibit highly distinct projection patterns. L5 ET cells project predominantly to subcortical targets while having limited intracortical collateral projections. L2/3/4 IT and L6 Car3 cortical/claustral neurons both have predominantly intracortical projections, but are easily distinguishable by the much more extensive and diffused intracortical projection and the lack of striatal projection from the latter.

Third, further morphological diversities exist within a major cell class or type. Both core- and matrix-type thalamic neurons exhibit strong nucleus-specific projection patterns, sometimes even allowing further subdivision of a nucleus (e.g. LP and MD). In the cortex, we were able to cluster both L5 ET and L6 Car3 neurons into multiple subtypes with robustly distinct projection patterns that are often correlated with the regions their somas reside in. Furthermore, we identified cell populations with additional unique projection targets. For example, L5 ET cells in the motor cortex have a highly distinct subtype that situates in deep L5 and projects to the medulla; a unique set of L6 Car3 cells in TEa/ECT and claustrum send projections to amygdala areas. L2/3/4 IT neurons exhibit largely continuous variation in their projection patterns, but still show regional differences. L4 IT cells in SSp mostly do not have long projections, a dramatic difference from L4 IT cells we found in MOp, MOs and SSs which all have long intracortical projections. Among the L2/3/4 IT neurons with long projections, those from the motor cortex have longer total axon arborizations than those from the somatosensory cortex.

Fourth, hierarchical organization of thalamocortical and corticocortical connections has been shown with mesoscale dataset based on feedforward and feedback connections ^57^. In this study, we further demonstrate hierarchical organization exists at single neuron level. The core-type thalamic neurons predominantly terminate in L4 and lower L2/3 of their cortical target areas, characteristic of feedforward projections. Most of our reconstructed matrix-type thalamic neurons have substantial axon branches in L1 in their cortical target areas, characteristic of feedback projections. In the cortex, we find that axon arbors of SSp neurons predominantly terminate in middle layers in all target areas, consistent with the low hierarchical position of SSp with all of its output pathways being feedforward. On the other hand, MOp neurons appear to show both feedforward, e.g. mid-layer projection to MOs, and feedback, e.g. L1 projection to SSp and SSs, patterns.

These wide range of projection patterns and rules revealed in our study suggest highly distinct functional roles between different major neuron classes and types, as well as a diverse range of potential ways for individual neurons to participate in circuit activities.

To bring all these findings into the general framework of cell type characterization and classification, a major remaining question is how the morphological and projectional diversities compare and correlate with the neurons’ molecular identities. We attempted to address this question with two approaches, using validated driver lines to define the major cell class or type level identities of reconstructed neurons and Retro-seq to obtain transcriptomic profiles of neurons projecting to specific targets. Both approaches show that major cell classes or types of neurons have highly distinct morphological and projection patterns, however, at more fine-grained levels within these major types, especially for cortical and claustral neurons, we also observe extensive morphological and projection diversities that cannot be accounted for by preexisting transcriptomic subtypes or clusters. For example, Retro-seq cells with highly distinct targets (e.g. SSp L2/3/4 IT cells projecting to MOp or SSs, L6 Car3 and claustral neurons belonging to distinct morphological clusters) were mapped to the same transcriptomic clusters. Previous studies showed that L2/3 SSp pyramidal neurons projecting to MOp or SSs have distinct intrinsic and network physiological properties ^20, 73^. Even though they may not belong to distinct transcriptomic subtypes, it will be interesting to find out if there are gene expression differences that correspond to the differential connectional and physiological properties for these neurons, as did for primary visual cortical neurons projecting differentially to medial or lateral higher visual areas ^74^.

The large diversity of axonal morphologies and projection patterns observed from genetically well-defined neuronal populations is striking but also consistent with an increasing body of related work ^22, 36^. The large number of target regions each neuron innervates and their elaborate pattern differences (e.g. layer distributions) would make it extremely challenging if not impossible to accurately discern the difference of projection patterns between individual neurons using other approaches including multiplexed retrograde tracing or tissue dissection of barcode-labeled brains. It underscores the necessity of scaling up the full neuronal morphology characterization effort to gain a true understanding of the extent of such diversity. Such knowledge is foundational for the understanding of brain connectivity and function.

The apparent lack of correlation between transcriptomic and morphological types in many cases is intriguing. It is possible that the current unsupervised clustering approach is insufficient to uncover the genes specifically relevant to morphology from the thousands of genes expressed by the neurons. Alternatively, it is possible that morphological/connectional specificity is established during circuit development and that the associated gene signatures exist only at that time ^75^.

In either case, the result emphasizes the importance of performing single cell characterization within multiple modalities and taking an integrated approach to describe and classify cell types in an unbiased and comprehensive manner. In the future, it will be important to develop methods that allow full morphology reconstruction and gene expression profiling to be conducted in the same cell, and apply them to the study of single cells in both adult stage and during brain development, so that potential molecular correlates of morphological/connectional features can be identified. This and other approaches together will ultimately lead to an integrated understanding of the extraordinary cellular diversity of the brain that underlies its function.

## ACKNOWLEDGMENTS

We are grateful to the In Vivo Sciences, Molecular Biology, Histology, and Imaging teams at the Allen Institute for their technical support. This work was supported by the Allen Institute for Brain Science and by multiple grant awards from institutes under the National Institutes of Health (NIH), including award number R01EY023173 from The National Eye Institute to H.Z., U01MH105982 from the National Institute of Mental Health and Eunice Kennedy Shriver National Institute Of Child Health & Human Development to H.Z., and U19MH114830 from the National Institute Of Mental Health to H.Z. and Q.L. Generation of TIGRE-MORF mouse line was partly funded by U01MH106008 and U01MH117079 from the National Institute Of Mental Health to X.W.Y. The content is solely the responsibility of the authors and does not necessarily represent the official views of NIH and its subsidiary institutes. This work is also funded by an initiative of Southeast University (SEU) to support Open Science oriented international collaboration. SEU supported the informatics data management and analysis pipeline at the SEU-Allen Joint Center. The work was also funded by the National Science Foundation of China (NSFC) grant 61721092 to Q.L., NSFC grant 61871411 to L.Q., the University Synergy Innovation Program of Anhui Province GXXT-2019-008 to L.Q., NSFC grant 91632201 to W.X., and by the Tiny Blue Dot Foundation to C.K. The Wenzhou Medical University reconstruction team received funding from the EPFL - Blue Brain Project. The authors wish to thank the Allen Institute founder, Paul G. Allen, for his vision, encouragement, and support.

## AUTHOR CONTRIBUTIONS

H.Z. conceptualized the study. H.P. envisioned and led the development of the computational and data analysis platform. M.B.V., T.L.D., B.T. and X.W.Y. generated the TIGRE-MORF (Ai166) mouse line. Z.J.H. provided Fezf2-CreER, Plxnd1-CreER and Tle4-CreER mouse lines. K.E.H., R.L. T.L.D., B.T. and J.A.H. contributed to the generation and characterization of specific transgenic mouse lines. H.G., A.L., S.Z., X.L., J.Y. and Q.L. conducted fMOST imaging. W.W., S.J., Y.Y. and C.H. handled the imaging data. L.Q., L.N. and H.P. developed methods for registration of fMOST datasets to CCF. Zhi Z., S.J., Y.Y., Yimin W. and H.P. developed software tools for data conversion and morphology reconstruction. Yun W., X.K., Y.L., L.L., P.L., Y.S., L.Y., S.Z., A.F., E.S., J.P., J. Y., G.H., A.L., and Z.D. performed manual and semi-automatic morphology reconstruction. L.L. co-developed the neuronal reconstruction pipeline at SEU-ALLEN and contributed to analysis of striatal and thalamic cell types. Yun W., P.X., J.A.H. and H.P. performed manual or computational classification of morphological types. Zhi Z., S.K. Zi Z., and S.A.S. assisted with morphological analysis. L.D. contributed to single neuron and population level projection analysis. Y.Z., D.L., and Yun W. collaborated with H.P. on developing a quality control method for neuron reconstruction. H.P., W.X., Z.G. and H.Z. collaborated in setting up the SEU-ALLEN data-production team. K.E.H., Q.W. and J.A.H. conducted anterograde AAV tracing. T.N.N. performed retrograde tracing. Z.Y., T.N.N. and B.T. conducted scRNA-seq data generation and analysis. S.M. and S.M.S. provided project management. L.E., M.J.H., B.T., L.N., S.A.S., J.A.H., H.G., Q.L., H.P., H.Z. and C.K. provided scientific management. H.Z. led the writing of the manuscript in consultation with all authors.

## METHODS

### Animal care and use

Both male and female transgenic mice ≥ P56 were utilized for all experiments. All animals were housed 3-5 per cage and maintained on a 12-hour light/dark cycle, in a humidity- and temperature-controlled room with water and food available *ad libitum*. All experimental procedures related to the use of mice were conducted with approved protocols in accordance with NIH guidelines, and were approved by the Institutional Animal Care and Use Committee (IACUC) of the Allen Institute for Brain Science.

### Transgenic mice

All transgenic crosses are listed in **Supplementary Table 1**. Data for systematic characterization of the expression pattern of each transgenic mouse line can be found in the AIBS Transgenic Characterization database (http://connectivity.brain-map.org/transgenic/search/basic).

Induction of CreERT2 driver lines was done by administration via oral gavage (PO) of tamoxifen (50 mg/ml in corn oil) at original (0.2 mg/g body weight) or reduced dose for one day in an adult mouse. The dosage for mice age P7-P15 is 0.04 ml. Mice can be used for experiments at 2 or more weeks after tamoxifen dosing. Specific dose of tamoxifen to induce sparse labeling in each CreERT2 driver line is shown in **Supplementary Table 1**.

### TissueCyte STPT imaging

Imaging by serial two-photon (STP) tomography (TissueCyte 1000, TissueVision Inc. Somerville, MA) has been described in earlier published studies ^16, 76^.

Mice were deeply anesthetized with 5% isoflurane and intracardially perfused with 10 ml of saline (0.9% NaCl) followed by 50 ml of freshly prepared 4% paraformaldehyde (PFA) at a flow rate of 9 ml/min. Brains were dissected and post-fixed in 4% PFA at room temperature for 3–6 h and then overnight at 4 °C. Brains were rinsed briefly with PBS and stored in PBS with 0.1% sodium azide until imaging.

Prior to imaging, the brain was embedded in a 4.5% oxidized (10 mM NaIO4) agarose solution in a grid-lined embedding mold to standardize its placement in an aligned coordinate space. The agarose block was then left at room temperature for 20 min to allow solidification. Cross-linking between brain tissue and agarose was promoted by placing the solidified block in 0.5% sodium borohydride in 0.5 M sodium borate buffer (pH 9.0) overnight at 4 °C. The agarose block was then mounted on a 1 × 3 glass slide using Loctite 404 glue and prepared immediately for serial imaging.

Image acquisition was accomplished using TissueCyte 1000 systems (TissueVision, Cambridge, MA) coupled with Mai Tai HP DeepSee lasers (Spectra Physics, Santa Clara, CA). The mounted specimen was fixed through a magnet to the metal plate in the centre of the cutting bath filled with degassed, room-temperature PBS with 0.1% sodium azide. A new blade was used for each brain on the vibratome and aligned to be parallel to the leading edge of the specimen block. Brains were imaged from the caudal end. The specimen was illuminated with 925 nm wavelength light through a Zeiss 320 water immersion objective (NA = 1.0), with 250 mW light power at objective. The two-photon images for red, green and blue channels were taken at 75 µm below the cutting surface. To scan a full tissue section, individual tile images were acquired, and the entire stage was moved between each tile. After an entire section was imaged, the x and y stages moved the specimen to the vibratome, which cut a 100-µm section, and returned the specimen to the objective for imaging of the next plane. The blade vibrated at 60 Hz and the stage moved towards the blade at 0.5 mm per sec during cutting. Images from 140 sections were collected to cover the full range of mouse brain at an x-y resolution of 0.35 µm per pixel. Upon completion of imaging, sections were retrieved from the cutting bath and stored in PBS with 0.1% sodium azide at 4°C.

### fMOST imaging

In summary, a GFP-labeled brain is first embedded in resin. The resin-embedded GFP fluorescence can be recovered through chemical reactivation ^77^ provided by adding Na_2_CO_3_ in the imaging water bath. Thus, a line-scanning block-face imaging system can be employed to maximize imaging speed. Following imaging of the entire block-face, the top 1-µm tissue is sliced off by a diamond knife, exposing the next face of the block for imaging. For the entire mouse brain, a 15-20 TB dataset containing ∼10,000 coronal planes of 0.2-0.3 µm X-Y resolution and 1 µm Z sampling rate is generated within 2 weeks.

All tissue preparation has been described previously ^78^. Following fixation, each intact brain was rinsed three times (6 h for two washes and 12 h for the third wash) at 4°C in a 0.01 M PBS solution (Sigma-Aldrich Inc., St. Louis, US). Then the brain was subsequently dehydrated via immersion in a graded series of ethanol mixtures (50%, 70%, and 95% (vol/vol) ethanol solutions in distilled water) and the absolute ethanol solution three times for 2 h each at 4°C. After dehydration, the whole brain was impregnated with Lowicryl HM20 Resin Kits (Electron Microscopy Sciences, cat.no. 14340) by sequential immersions in 50, 75, 100 and 100% embedding medium in ethanol, 2 h each for the first three solutions and 72 h for the final solution. Finally, each whole brain was embedded in a gelatin capsule that had been filled with HM20 and polymerized at 50°C for 24 h.

The whole brain imaging is realized using a fluorescence micro-optical sectioning tomography (fMOST) system. The basic structure of the imaging system is the combination of a line-scanning upright epi-fluorescence microscopy with a mechanic sectioning system. This system runs in a line-scanning block-face mode but updated with a new principle to get better image contrast and speed and thus enables high throughput imaging of the fluorescence protein labeled sample (manuscript in preparation). Each time we do a block-face fluorescence imaging across the whole coronal plane (X-Y axes), then remove the top layer (Z axis) by a diamond knife, and then expose next layer, and image again. The thickness of each layer is 1.0 µm. In each layer imaging, we used a strip scanning (X axis) model combined with a montage in Y axis to cover the whole coronal plane ^79^. The fluorescence, collected using a microscope objective, passes a bandpass filter and is recorded with a TDI-CCD camera. We repeat these procedures across the whole sample volume to get the required dataset.

The objective used is 40X WI with numerical aperture (NA) 0.8 to provide a designed optical resolution (at 520 nm) of 0.35 μm in XY axes. The imaging gives a sample voxel of 0.35 x 0.35 x 1.0 μm to provide proper resolution to trace the neural process. The voxel size may vary for difference objectives. Other imaging parameters for GFP imaging include an excitation wavelength of 488 nm, and emission filter with passing band 510-550 nm. The fMOST is a two-color imaging system. The green channel is used to obtain the complete morphology of neurons, and the red channel is used to obtain the cellular architecture information of propidium iodide(PI) staining.

### Full neuronal morphology reconstruction system

We usedVaa3D, an open-source, cross-platform visualization and analysis system, for the tasks of reconstructing massive neuronal morphologies. To efficiently and effectively deal with the whole-mouse brain imaging data, we incorporated several enabling modules into Vaa3D, such as TeraFly, TeraVR, and a number of other supporting tools. TeraFly supports visualization and annotation of multidimensional imaging data with virtually unlimited scales. A user can flexibly choose to work at a specific region of interest (ROI) with desired level of detail (LoD). TeraVR is an annotation tool for immersive neuron reconstruction that has been proved to be critical for achieving precision and efficiency in morphology data production. It creates stereo visualization for image volumes and reconstructions and offers an intuitive interface for the user to interact with such data. Both TeraFly and TeraVR are seamlessly integrated in Vaa3D and can be used combinedly and flexibly. From reconstructions (in SWC file format), morphological quantification statistics is obtained to characterize neurons. QC process identifies errors based on morphological indicators and does corrections in a feedback setting. QC process then refines the skeleton location with Mean-Shift ^80^ and performs pruning focused on terminal location refinement. When needed auto-refinement fits the tracing to the center of fluorescent signals. The whole process ends with SWC resampling and registration. The final reconstruction of each neuron is a valid single tree without breaks, loops, multiple branches from a single point, etc.

### Registration to CCF

We used mBrainAligner based on BrainAligner ^81^ to perform 3D registration from fMOST images (subject) to the average mouse brain template of CCFv3 (target) (Extended Data Fig. 3). The main steps are: 1) fMOST images were first down-sampled by 64×64×16 (X, Y, Z) to roughly match the size of target brain. 2) The stripe artifacts in fMOST images that raised from diamond knife cutting and imaging process were eliminated by using log-space frequency notch filter. 3) The dense outer-contour feature points of target and subject brain (about 1500 points per brain) were uniformly sampled from the brains’ outer-contour that obtained using adaptive threshold, and then affine aligned using a reliable landmark points matching algorithm to ensure the subject brain has the same position, orientation and scale as the target brain. 4) Intensity was normalized by matching the local average intensity of subject image to that of target image in a sliding window manner with patch size 41×41×41 and stride one. 5) For the target brain, 1,744 landmarks corresponding to the points of high curvature (corners or junction of different brain compartments) in CCFv3 annotation image were detected via 3D Harris corner detector. Based on a combination of texture, shape context and deep-learning-derived features, mBrainAligner established the correspondence between target and subject brain by iteratively deforming these target landmarks to fit the subject image, and accomplished the local alignment using the smooth-thin-plate-spline (STPS). 6) Finally, to ensure the accuracy of registration, automatic registration results were examined in the semi-automatic registration module of mBrainAligner, and if necessary, the boundaries of brain region were further optimized in a manual or semi-automatic way. Once images were aligned, the reconstructed neurons and somas were warped to CCF space using the generated deformation fields.

### Processing single cell morphological data

Pre-processing of SWC files: SWC files were processed and examined with Vaa3D plugins to ensure topological correctness: sorted single tree with root node as soma. Terminal branches < 10 pixels were pruned to remove artifacts. SWC files were resampled with a step size of 64 (x), 64 (y) and 16(z) before registration.

Quantification of axon projection patterns: To analyze the distribution and amount of axon in brain-wide targets following registration to the CCFv3, we used a manually curated set of 316 non-overlapping structures at a mid-ontology level that are most closely matched in size or division. Ipsi- and contra-lateral sides of brain regions were calculated separately.

Morphological features: Axonal and dendritic morphological features, defined according to L-measurement (Scorcioni et al., 2008), were calculated using Vaa3D plugin “global_neuron_feature”. Selected features include:

Axon global: ‘Overall Width’, ‘Overall Height’, ‘Overall Depth’, ‘Total Length’, ‘Euclidean Distance’, ‘Max Path Distance’, ‘Number of Branches’.

Axon local: ‘Total Length’, ‘Number of Branches’.

Dendrite: ‘Overall Width’, ‘Overall Height’, ‘Overall Depth’, ‘Total Length’, ‘Max Euclidean Distance’, ‘Max Path Distance’, ‘Number of Branches’, ‘Max Branch Order’.

Local axons were defined as axon arbors within 200 microns from the somata. Local axons and dendrites were rotated based on principle component analysis (PCA) so dimensions were aligned with the largest to smallest spans. Then shifting was performed to localize somata at the origin of coordinates.

### mBrainAnalyzer

Our mBrainAnalyzer toolbox, which was developed for analysis of full neuron morphology, includes multiple modules for feature quantification, arbor detection, statistical analysis and visualization. In addition to morphological features (e.g. total length, angle of branches etc.), this toolbox also quantifies projection intensities at branch length level and number of terminal levels. Using the arbor detection module, one can define sub-cellular components of a neuron as the granularity. Analysis and visualization can be performed at both whole-cell and arbor levels.

### Arbor detection and partition

We detected and partitioned a series of neuronal arbors out of each neuron reconstruction using a graph-partition clustering method. First, as a neuron consists of a number of topologically connected reconstruction nodes, the neuron was viewed as a graph, where every reconstruction node (unit) in the neuron was connected with its parent node with an edge specified by the topological connection of the parent-child pair with the edge weight, or ‘similarity’ *s*, set to be the exponential of the negative 3D Euclidean distance, *d*, of these two nodes, i.e. *s* = exp(-*d*). Then, we considered the normalized graph-cut method ^82^ to extract “clusters” of reconstruction nodes so that the within-cluster “total similarity” of nodes would be maximized and cross-cluster total similarity would be minimized. As a result, each such coherent cluster corresponds to one neuron arbor, which was also visually checked to ensure its correctness. Third, to automatically determine the number of such clusters, for a presumed number of clusters, we calculated the normalized score of total cross-cluster similarity divided by the total within-cluster similarity, followed by trial-testing a range (between 2 to 8) of such presumed cluster-numbers to determine the optimal number that would minimizes this normalized score. In the final result, the detected arbor that contains the soma is called soma-arbor; the remaining arbors are called non-soma arbors.

### Feature quantification of cortical arbors

We divided the cortex into consecutive coronal slices of 100μm thick. Anchor points were evenly sampled along the outer border of each slice, with normal vectors that are perpendicular to the local cortical surface and pointing to the inside of the brain. Nodes of arbors were assigned to their neighbor anchors and projected onto the surface by corresponding normal vectors. Depth of nodes were determined by the length of projection along normal vectors. We also estimated the area of an anchor by their distance to neighbor anchors and slice thickness. The 2D cortical area of an arbor was determined by the total areas of unique anchors occupied by its nodes. To determine the radius of an arbor, we assigned arbor ‘center’ as the node that has the shortest average distance to other nodes. Radius was determined by a growing sphere until 70% segments are inside it. For neurons with tufted apical dendrite, we vertically shifted the arbors, so the top of apical dendrites reached L1. We manually confirmed that all tufted apical dendrites reached L1 in the original image.

### Clustering of cortical arbors

For local (soma-neighboring) arbors, the following features were used for clustering: ‘2d_area’, ‘axon_length’, ‘dend_length’, ‘radius’, ‘axon_depth_mean’, ‘axon_depth_std’, ‘dend_depth_mean’, ‘dend_depth_std’. Axon/Dendrite depth features were normalized by the average thickness of cortical areas where the arbor locates. We performed PCA to reduce the effect of noise. Top principle components were selected to recover 95% of variance. We applied UMAP dimension reduction using python package ‘UMAP’ (McInnes et al., 2018). The ‘n_neighbors’ parameter was set at 4. K-means clustering was performed using the UMAP embeddings as input.

For distal arbors, we profiled the axon density distribution along cortical depth as input features of clustering. We did not use the same features as local arbors as distal arbors exhibited much higher diversity. Clustering approach for distal arbors is the same as local arbors.

### Neuron-beta

we developed the Neuron-beta metric by borrowing the concept of the ‘Beta’ value from the finance field ^83^. For each group, defined by brain areas and/or cortical layers, we calculated the average of mesoscale experiments as *M* = [*m_1_, …, m_p_*], *p* = number of brain areas. For one single cell *S* = [*s_1_, …, s_p_*], we define the neuron-beta value as:

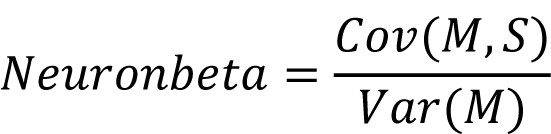

### Clustering of cortical L6 Car3 and claustral neurons

Data normalization: Morphological features were normalized by the mean and standard variation in a feature-wise manner. Projection pattern features were normalized by the total length per 100 µm in a sample-wise manner and scaled by logarithm. Soma locations were flipped to the same hemisphere.

Similarity metrics: For each feature set, we first calculated the Euclidean distance matrix. Then a ranked K-nearest neighbor (KNN) matrix was created. We then applied the Shared Nearest Neighbor (SNN) approach to measure the similarity between each pair of samples *x_i_* and *x_j_*. The SNN metric was defined as the maximum average rank among their common neighbors:

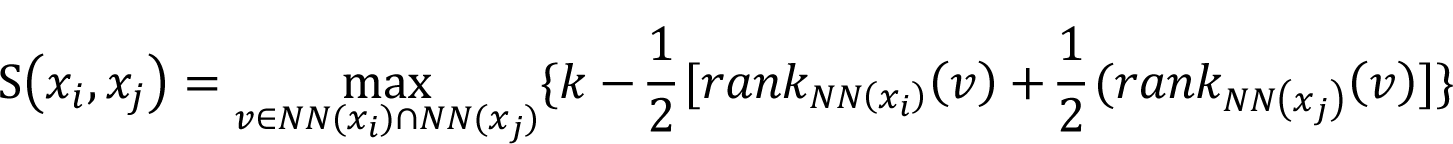

Similarity scores were set as 0 for pairs with non-overlapping KNN sets and a weighted SNN graph is created.

Co-clustering analysis: Co-clustering matrix for each feature set was calculated by iterative random sampling. During each iteration, 95% of samples were randomly selected to create an SNN graph. We then applied the “Fast-greedy” community detection algorithm using python package “python-igraph” for clustering assignment. For each pair of samples, the co-clustering score was defined as the times of co-clustering normalized by the iterations of co-occurring. Resampling was performed 5,000 times to reach saturation. The overall co-clustering matrix is a weighted average of the four feature sets. Agglomerative clustering was performed to the co-clustering matrix to get clusters.

Outlier removal: Outliers were detected by comparing the Euclidean distance between a sample and the other samples with the same cluster identity. We used overall within-cluster distance as the background distribution. Samples with significantly higher (one-sided Mann-Whitney test) within-cluster distance were filtered out as outliers. Agglomerative clustering was performed for the remaining co-clustering matrix. This process iterated until no new outlier could be detected.

Characterization of cell types: For each feature set, we performed two-sided Mann-Whitney tests: claustrum vs. cortical neurons; each cluster vs. other clusters. P-values were adjusted by Bonferroni correction.

### Anterograde tracing and retrograde labeling

For anterograde projection mapping, we injected AAV2/1-pCAG-FLEX-EGFP-WPRE-pA (Oh et al., 2014) into CLA, SSs or SSp of Gnb4-IRES2-Cre or Gnb4-IRES2-CreERT2 mice at P37-P65 respectively. Stereotaxic injection procedures were performed as previous described ^16^ and stereotaxic coordinates used for each experiment can be found in the data portal. For the Gnb4-IRES2-CreERT2 mice, tamoxifen induction was conducted one week post injection at full dose (0.2 mg/g body weight) for 5 consecutive days. Mice survived 3 weeks (or 4 weeks for the tamoxifen-induced mice) post injection, and brains were perfused and collected for TissueCyte imaging.

For retrograde labeling, we injected several different types of retrograde viral tracers, including AAV2-retro-EF1a-dTomato or AAV2-retro-EF1a-Cre ^84^, RVdGdL-Cre or RVdL-FlpO ^85^, or CAV2-Cre ^86^, into specific target regions of defined transgenic mice (**Supplementary Table 4**). We FACS-sorted and collected RFP+ or RFP+/GFP+ cells from defined source regions for single-cell RNA-sequencing. Stereotaxic injection procedures were performed as previous described ^16^. Mice were injected at P40 or older, with 16-31 days survival post-injection.

### Single-cell RNA-sequencing

Cells from transgenic mice or transgenic mice injected with retrograde tracers were collected by microdissection of different cortical regions. Single-cell suspensions were created and cells were collected using fluorescence activated cell sorting (FACS). FACS gates were selective for cells with fluorescent protein expression from transgenic and/or viral reporters.

Cells were then frozen at −80°C, and were later processed for scRNA-seq using the SMART-Seq v4 method ^4^. After sequencing, raw data was quantified using STAR v2.5.3 and were aligned to both a Ref-Seq transcriptome index for the mm10 genome, and a custom index consisting of transgene sequences. PCR duplicates were masked and removed using STAR option ‘bamRemoveDuplicates’. Only uniquely aligned reads were used for gene quantification. Gene read counts were quantified using the summarizeOverlaps function from R GenomicAlignments package using both intronic and exonic reads, and QC was performed as described ^4^.

Clustering was performed using house developed R package scrattch.hicat (available via GitHub https://github.com/AllenInstitute/scrattch.hicat). In additional to classical single-cell clustering processing steps provided by other tools such as Seurat, this package features automatically iterative clustering by making finer and finer splits while ensuring all pairs of clusters, even at the finest level, are separable by fairly stringent differential gene expression criteria. The package also performs consensus clustering by repeating iterative clustering step on 80% subsampled set of cells 100 times, and derive the final clustering result based on cell-cell co-clustering probability matrix. This feature enables us to both fine tune clustering boundaries and to assess clustering uncertainty. One critical criterion that determines the clustering resolution is the minimal differential gene expression (DGE) requirement between all pairs of clusters. Using stringent DGE requirement results in fewer clusters with more prominent differences between clusters, while using more relaxed DGE expression result in more clusters captured by more subtle differences. For the whole cortical and hippocampal dataset with ∼75,000 cells, we used the standard DGE requirement as in ^4^. More specifically, q1.th = 0.5 (minimal fraction of cells in a given cluster that express the positive markers), q.diff.th=0.7 (normalized differences in fraction of cells expressing the positive markers between the foreground and background cluster, maximal value is 1), and de.score.th=150 (overall assessment of the statistical significance of all DGE genes).

### Data and Code availability

The raw and TeraFly converted fMOST image datasets of all mouse brains used in this study, as well as the CCFv3 registered single neuron reconstructions, are available at BICCN’s Brain Image Library (BIL) at Pittsburgh Supercomputing Center (www.brainimagelibrary.org). The single neuron reconstructions, the CCFv3 registered version of these reconstructions, as well as 3D navigation movie-gallery of these data are available at SEU-ALLEN Joint Center, Institute for Brain and Intelligence (https://braintell.org/projects/fullmorpho/). The neuron reconstructions are also released through NeuroMorpho.Org. The Vaa3D platform along with the TeraFly and TeraVR reconstruction software is available through the GitHub release page of vaa3d.org. The mBrainAligner and mBrainAnalyzer software packages are available upon request.

Mesoscale tracing data (including high resolution images, segmentation, registration to CCFv3, and automated quantification of injection size, location, and distribution across brain structures) are available through the Allen Mouse Brain Connectivity Atlas portal (http://connectivity.brain-map.org/). Retro-seq SMART-Seq v4 data have been deposited to BICCN’s NeMO Archive.

## Supplemental Information

**Supplementary Table 1**. Transgenic mice used for the generation of fMOST imaging datasets, including main metadata information and tamoxifen dosing (see Methods) for sparse labeling.

**Supplementary Table 2**. List of reconstructed neurons, with each neuron’s 3D coordinates, annotated soma location in CCFv3 after registration and manual correction, transgenic line and brain ID, neuron subclass or type assignment, and projection matrix.

**Supplementary Table 3**. Mesoscale anterograde tracing experiments used in this study for comparison with single neuron projection patterns, including main metadata information and projection matrix.

**Supplementary Table 4**. Retro-seq cells for scRNA-seq analysis, with relevant metadata including retrograde labeling information.

---

**Extended Data Figure 1.**
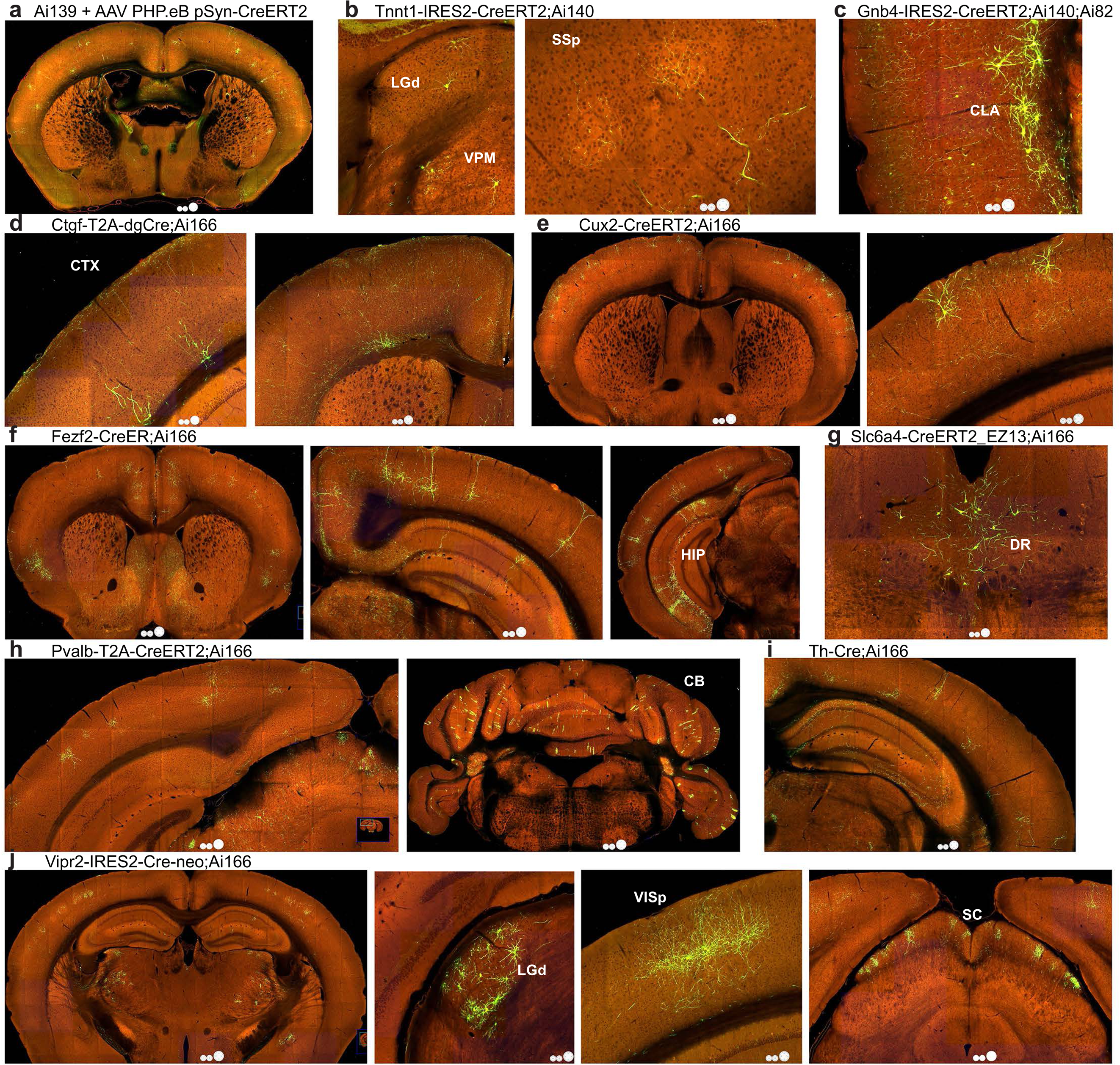
Representative TissueCyte images with sparse and strong labeling of various neuron types using these approaches. **a**, A Synapsin I promoter-driven CreERT2-expressing AAV serotyped with PHP.eB was delivered at a dilution of 1:1000 by retroorbital injection into an Ai139 mouse, followed by a 1-day tamoxifen induction one week post injection, resulting in random sparse labeling of neurons throughout the brain. **b**, In a Tnnt1-IRES2-CreERT2;Ai140 brain, low-dose tamoxifen induction results in sparse labeling of thalamic projection neurons (left panel) with their axon terminal clusters in cortex clearly visible (right panel). **c**, In a Gnb4-IRES2-CreERT2;Ai140;Ai82 brain, low-dose tamoxifen induction results in sparse labeling of *Gnb4*+ claustral and cortical neurons with their widely dispersed axon fibers clearly visible. **d**, Cortical L6b neurons in a Ctgf-T2A-dgCre;Ai166 brain. **e**, Cortical L2/3/4 neurons in a Cux2-CreERT2;Ai166 brain. **f**, Cortical L5 ET neurons in a Fezf2-CreER;Ai166 brain. **g**, Serotonergic neurons in dorsal raphe (DR) in a Slc6a4-CreERT2_EZ13;Ai166 brain. **h**, Interneurons in cortex and cerebellum in a Pvalb-T2A-CreERT2;Ai166 brain. **i**, *Th*+ cortical interneurons in a Th-Cre;Ai166 brain. **j**, Projection neurons in LGd and other thalamic nuclei in a Vipr2-IRES2-Cre-neo;Ai166 brain. Third panel, axon projections from LGd neurons are seen in primary visual cortex (VISp). Fourth panel, axon projections likely from retinal ganglion cells are seen in superior colliculus (SC). Tamoxifen doses for CreERT2-containing mice are shown in **Supplementary Table 1**.

**Extended Data Figure 2.**
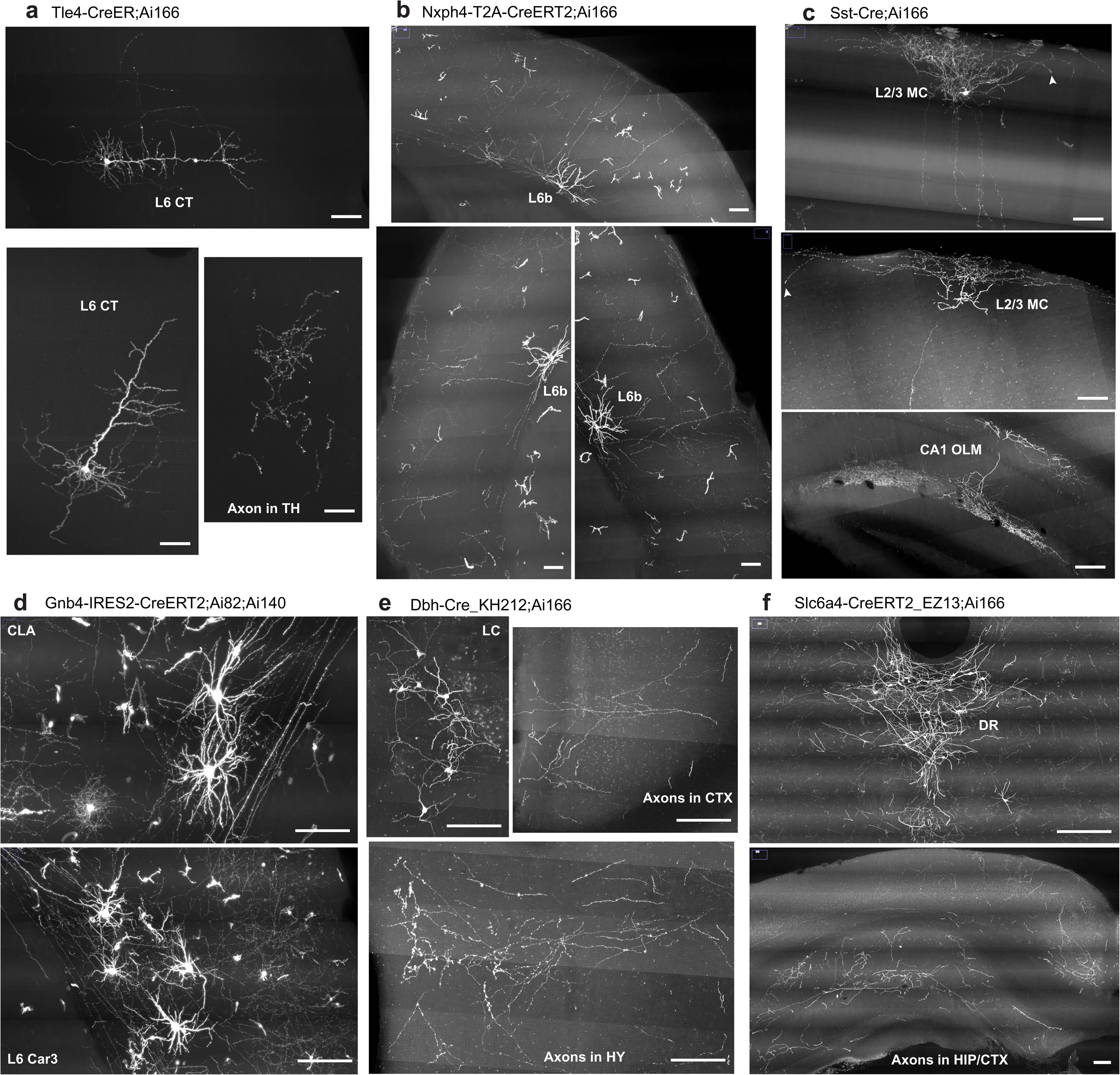
Sparse, robust and consistent labeling and visualization of the dendritic and axonal arborizations of neuronal types in additional Cre lines. **a,** Cortical L6 CT neurons and their characteristic apical dendrites not reaching L1, as well as local axon collaterals and long-range axon projections into thalamus (TH), labeled in a Tle4-CreER;Ai166 brain. **b,** Cortical L6b neurons and their local axon projections up into L1 seen in a Nxph4-T2A-CreERT2;Ai166 brain. **c,** Cortical inhibitory Martinotti cells (MC) and hippocampal CA1 OLM cells labeled in a Sst-Cre;Ai166 brain. **d,** *Gnb4*+ claustral (CLA) and cortical (L6PC) neurons with their widely dispersed axon fibers seen in a Gnb4-IRES2-CreERT2;Ai140;Ai82 brain. **e,** Noradrenergic neurons labeled in the locus ceruleus (LC), and their long-range axon fibers seen in cortex (CTX) and hypothalamus (HY) in a Dbh-Cre_KH212;Ai166 brain. **f,** Serotonergic neurons labeled in the dorsal raphe (DR), and their long-range axon fibers seen in hippocampus (HIP) and cortex (CTX) in a Slc6a4-CreERT2_EZ13;Ai166 brain. Images shown are 100-µm maximum intensity projection (MIP) images (*i.e.*, projected from 100 consecutive 1-µm image planes). Arrowheads indicate observed terminal boutons at the end of the axon segments. Tamoxifen doses are shown in **Supplementary Table 1**. Scale bars, 100 µm.

**Extended Data Figure 3.**
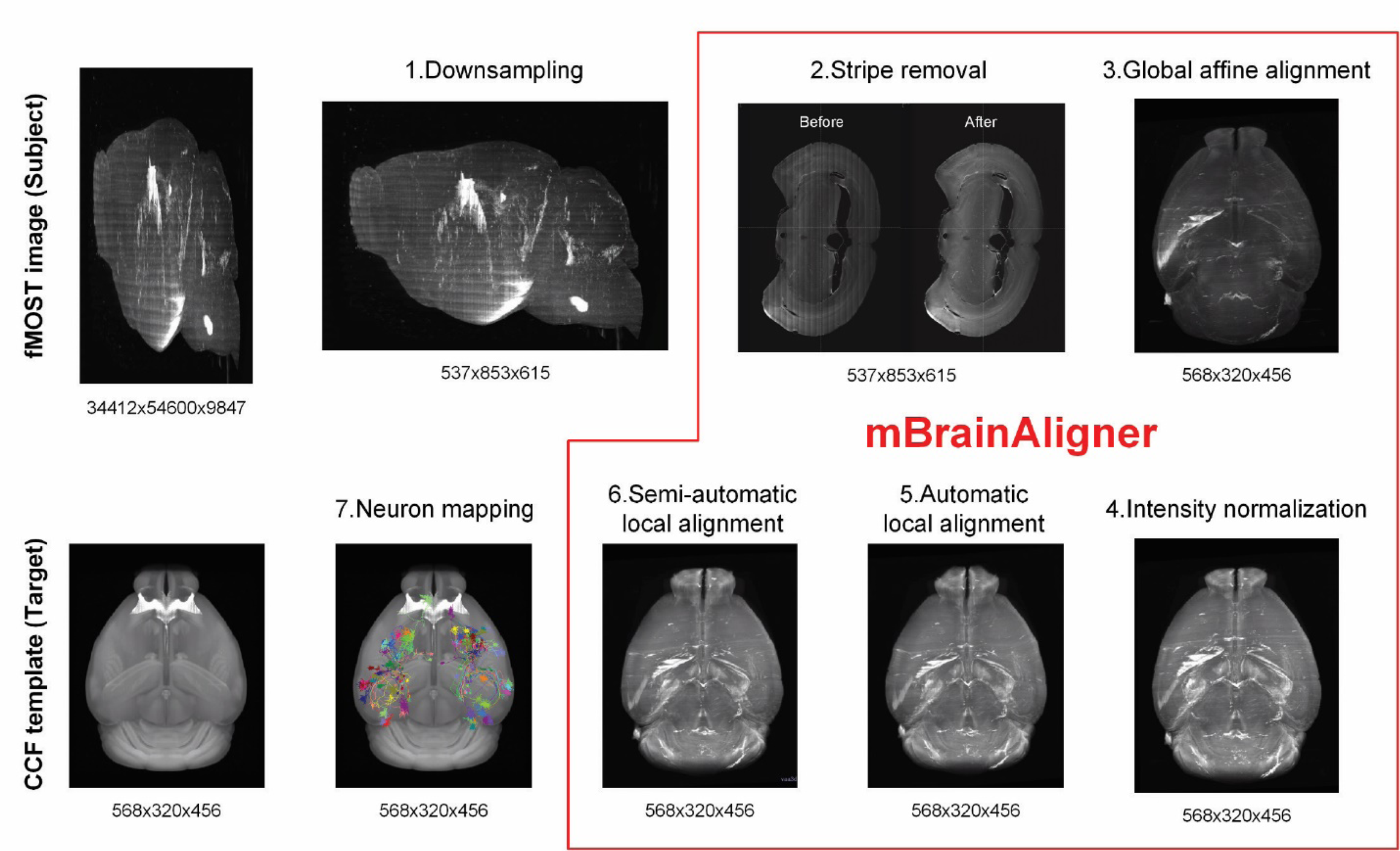
CCF registration workflow. Pipeline of 3D registration from fMOST image (subject) to average mouse brain template of CCFv3 (target). Numbers below each panel indicate the pixel sizes in the order of X*Y*Z. See Methods for explanation of each step.

**Extended Data Figure 4.**
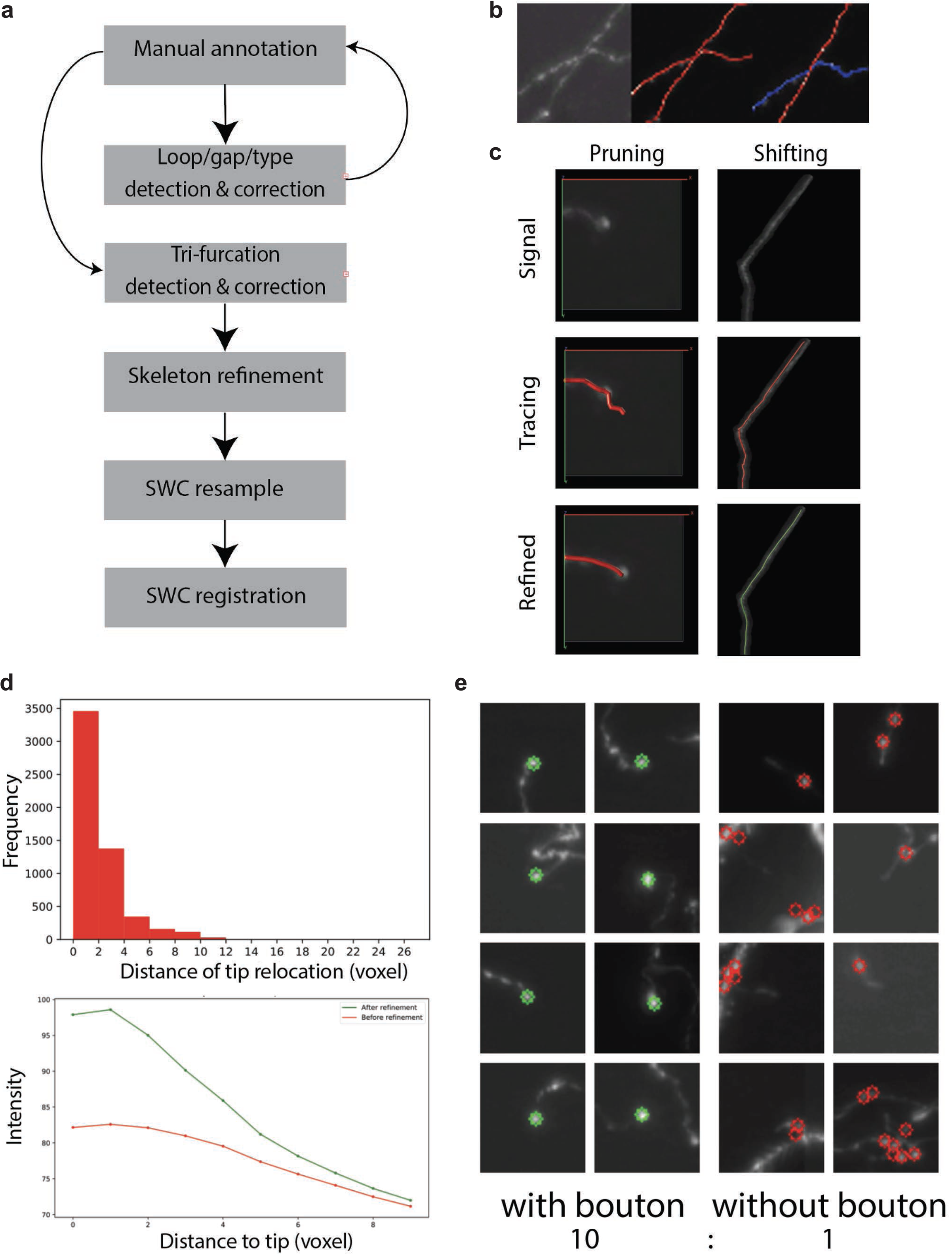
QC of reconstructed morphologies. **a,** Workflow of SWC post-processing process for QC: 1. automatic detection and correction of basic reconstruction errors including loops, gaps and incorrect node types. 2. Corrections are sent back to manual verification. 3. Automatic detection and correction of trifurcation, which are usually overlapping neurites, instead of branching points. 4. Refinement of SWC files, including pruning of over-traced terminals and shifting skeleton to fit the center of image signals. 5. Resampling of SWC to achieve evenly distributed nodes. 6. SWC registration to the standard CCFv3 mouse brain template. **b,** Examples of trifurcation before (middle) and after (right) correction. Blue and red branches do not cross. **c,** Examples of refinement before and after pruning (left) and shifting (right). **d,** Refinement leads to more precisely defined axon termination. Upper, distribution of terminal relocation distance by pruning. Lower, radius-decay curve of terminal signals shows that after refinement the axon ends at a brighter spot (indicating a bouton) rather than tapering off. **e,** Examples of axonal terminals that end with or without a bouton.

**Extended Data Figure 5.**
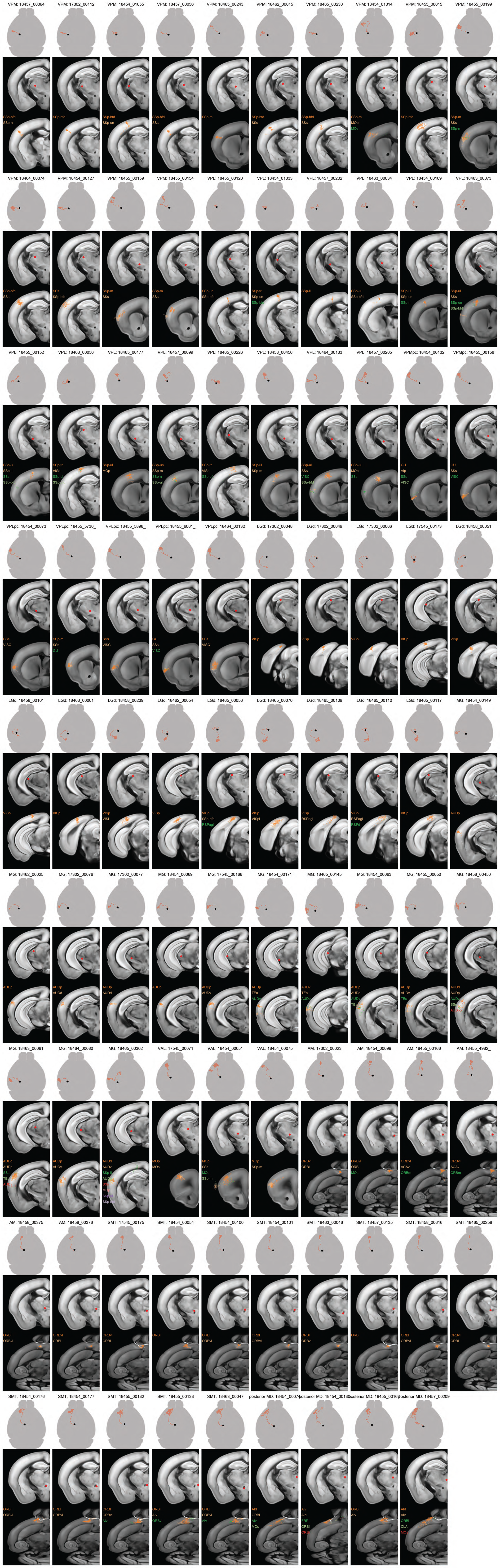
Tri-views of reconstructed neurons from “core” thalamic nuclei, visualized within the CCFv3 3D reference space. Each tri-view contains three views of the same neuron ordered from top to bottom: a whole-brain top-down view (soma indicated by a star, axon in red), a coronal plane showing the location of the soma (red dot), and a chosen coronal or horizontal plane close to the center of the main axon arbor with superimposed maximum projection view of the axon arbors. Cortical target regions with axon length >1 mm are indicated by different colors while other axon branches are shown in white.

**Extended Data Figure 6.**
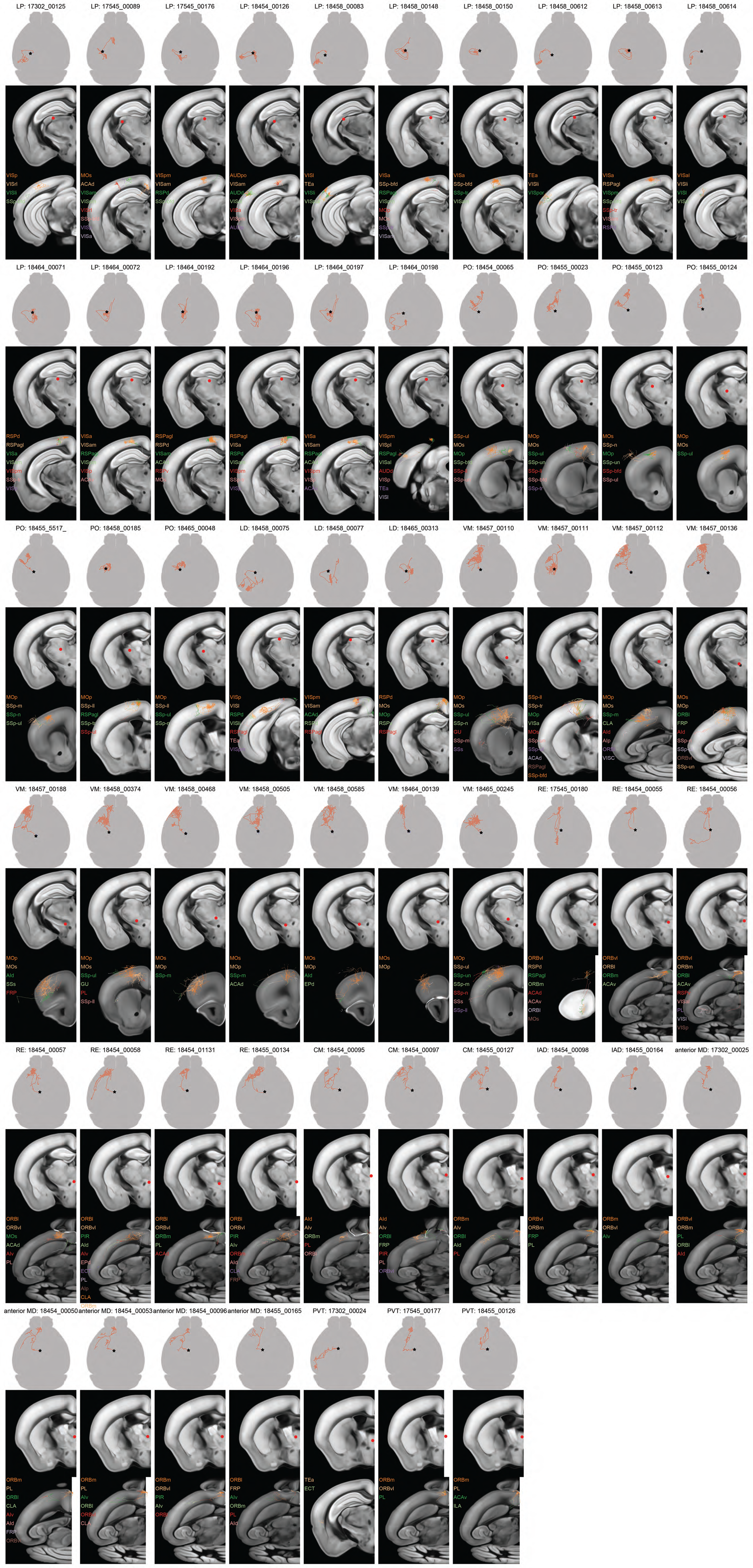
Tri-views of reconstructed neurons from “matrix” thalamic nuclei, visualized within the CCFv3 3D reference space. Each tri-view contains three views of the same neuron ordered from top to bottom: a whole-brain top-down view (soma indicated by a star, axon in red), a coronal plane showing the location of the soma (red dot), and a chosen coronal plane close to the center of the main axon arbor with superimposed maximum projection view of the axon arbors. Cortical target regions with axon length >1 mm are indicated by different colors while other axon branches are shown in white.

**Extended Data Figure 7.**
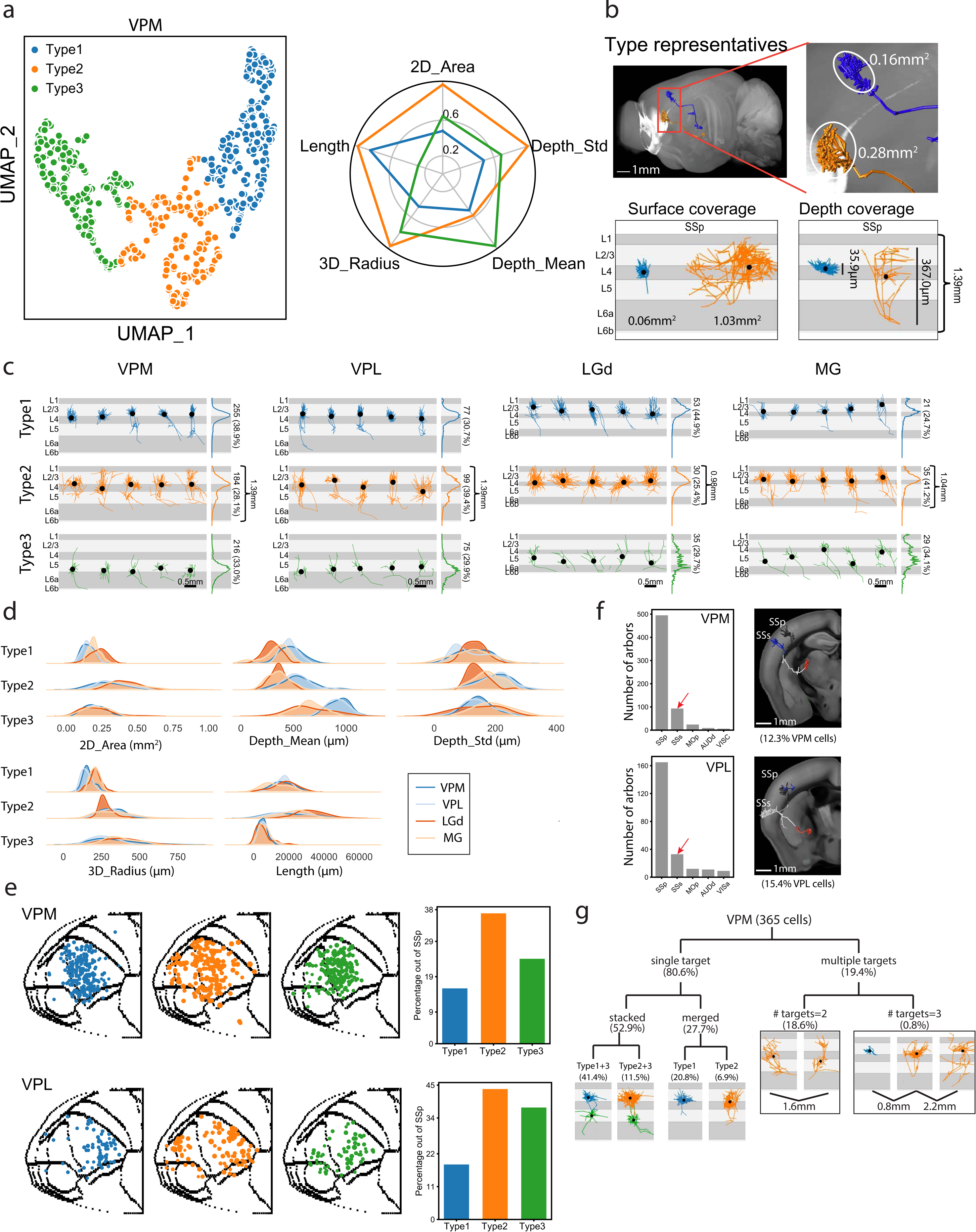
Thalamocortical axon arbor analysis. **a,** Clustering result indicates three types of cortical axon arbors in VPM neurons. Left, UMAP representation of VPM axon arbors, colored by cluster ID’s. Right, Polar plot of main features, values as normalized cluster averages. **b,** Representative (upper) and extreme (lower) examples of VPM cortical arbors. **c,** Examples grouped by thalamic nuclei and arbor types. In each sub-panel, vertical views are shown for 5 representative arbors, with branch length distribution for all neurons of the same cluster on the right side. Arbor numbers and percentage of the group are shown on the right side. **d**, Distribution of features grouped by thalamic nuclei and arbor types. **e**, Arbor locations of VPM and VPL neurons in 2D cortical map grouped by arbor types. Each dot represents the center of an arbor. Right panels show percentage of arbors outside of the primary target of VPM/VPL neurons. **f**, (Left) Counts of VPM/VPL arbors in cortical regions. (Right) Examples of neurons with double arbors, one in SSp and the other in SSs. **g**, Variation of VPM neurons by arbor compositions. ‘Single target’ neurons are described as ‘stacked’ or ‘merged’ by bi-layer or single-layer distribution. The stacked and merged groups can be further separated by arbor types. The ‘multiple targets’ group is divided by number of targets.

**Extended Data Figure 8.**
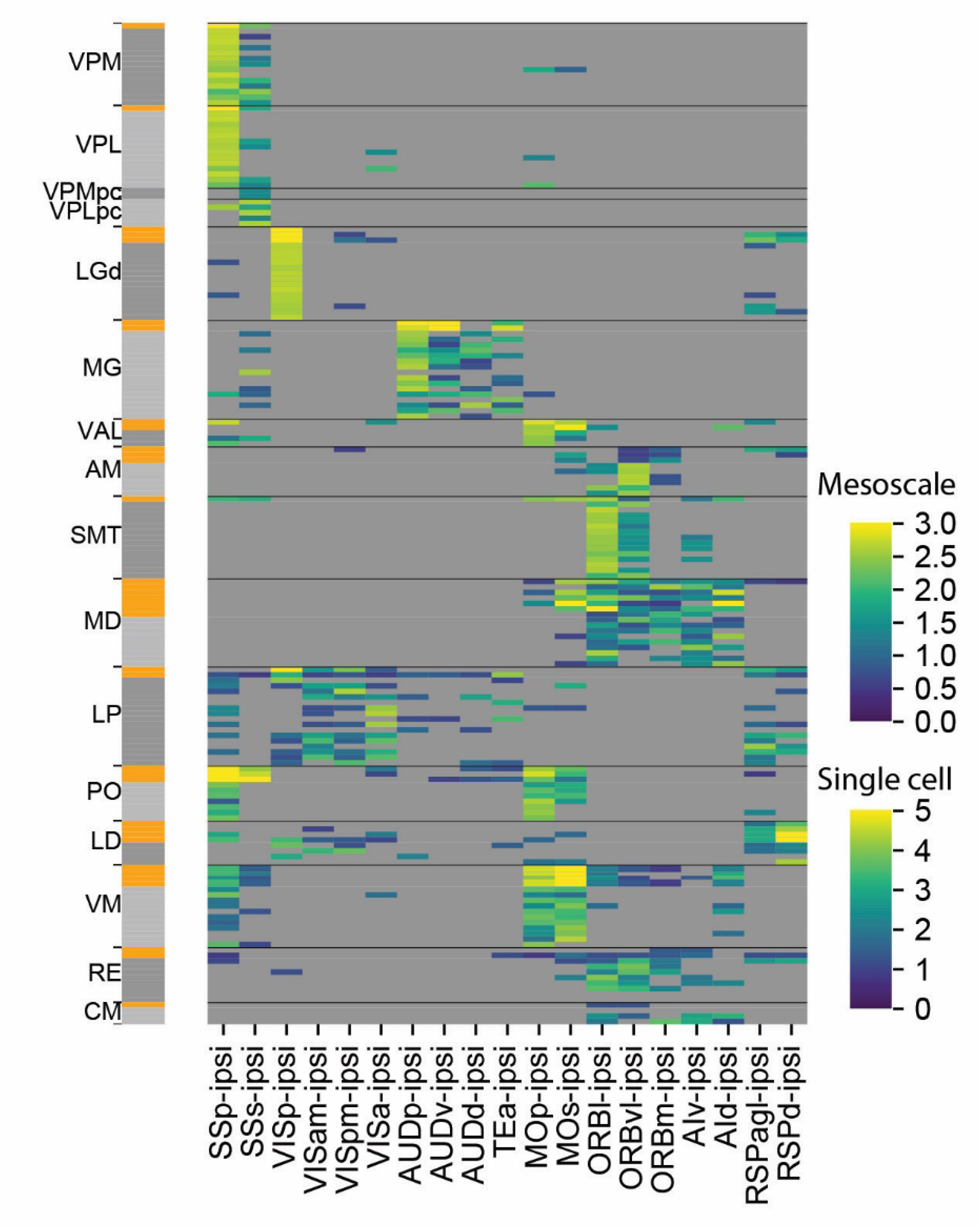
Comparison of thalamocortical projection patterns between mesoscale experiments and single neurons as well as among individual neurons, for each listed thalamic nucleus. Heatmap colors represent log (percentage of projection strength + 1), scaled to 0-3 for mesoscale experiments and 0-5 for single cell reconstructions. We set regions below a cutoff 0.5 as grey. The side bar to the left of heatmap indicates experiment types (single cell: grey, mesoscale: orange).

**Extended Data Figure 9.**
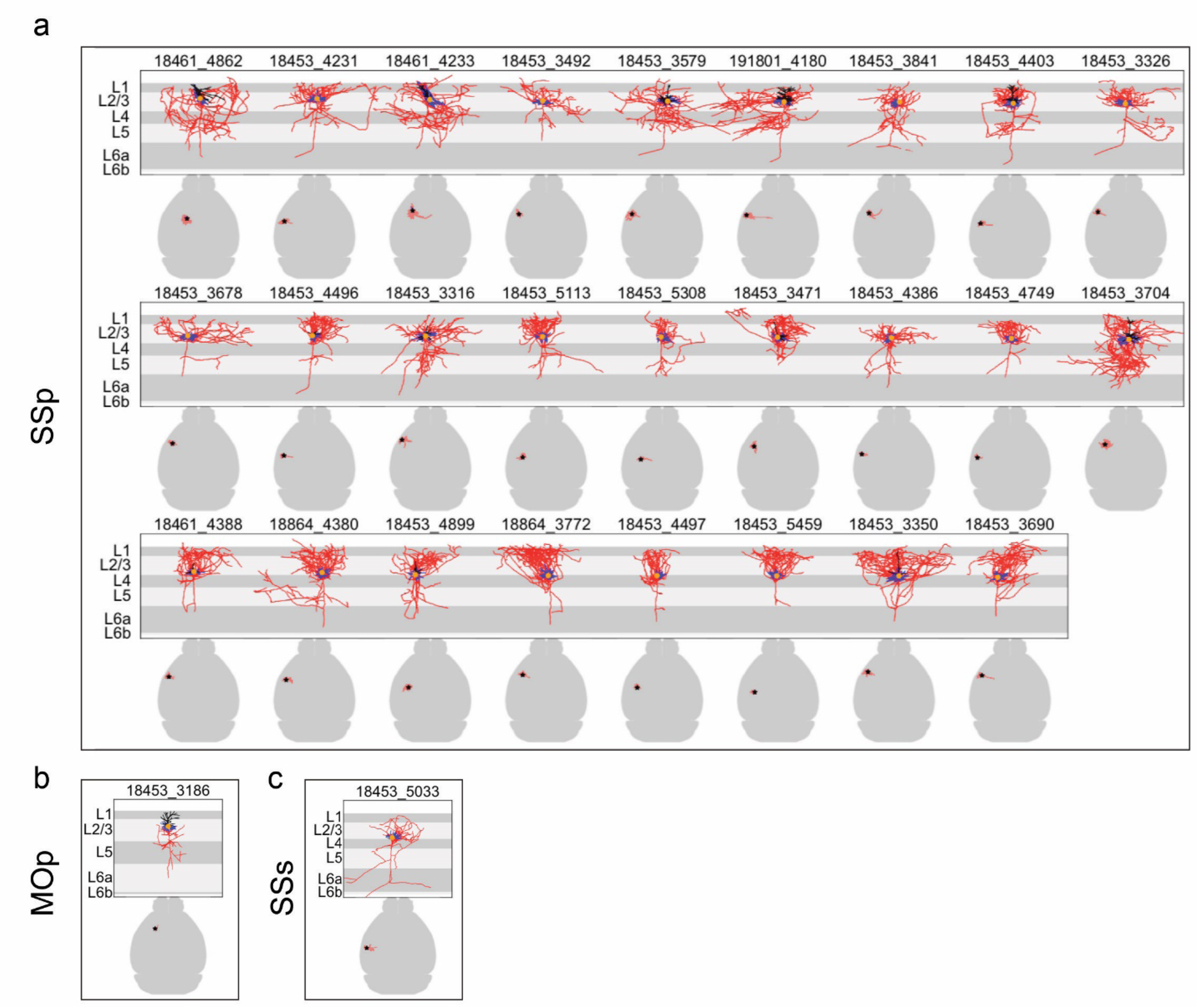
Reconstructed cortical L2/3 and L4 IT neurons of SSp, SSs and MOp without substantial long-range axon projections. For each neuron, both local morphologies (upper panels; apical dendrite in black, basal dendrite in blue, axon in red, soma as an orange dot) and whole-brain projections (lower panels; axon in red, soma as a star) are shown.

**Extended Data Figure 10.**
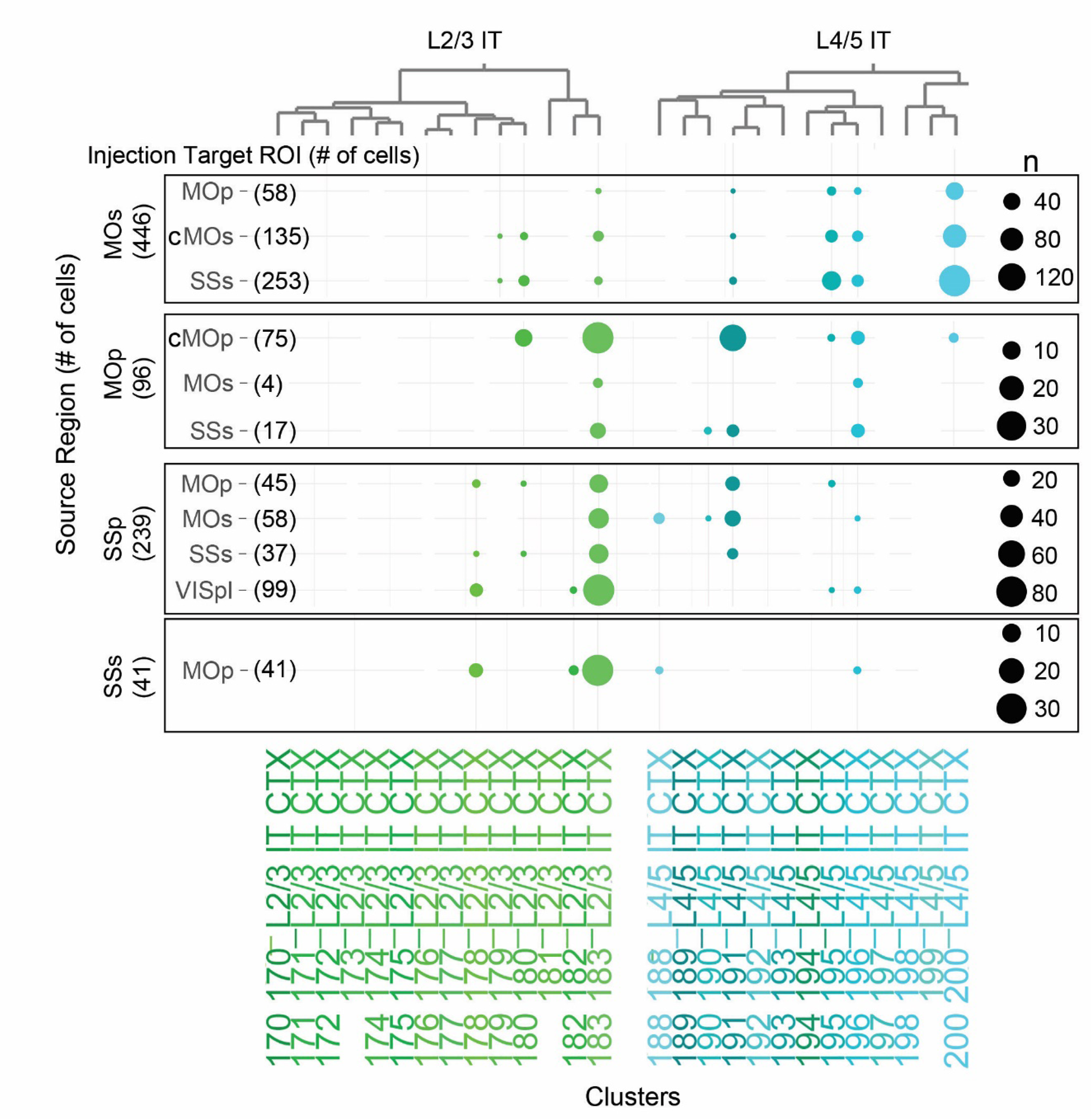
Retro-seq characterization of cortical L2/3 and L4/5 IT neurons from MOs, MOp, SSp and SSs. Transcriptomes of retrogradely labeled neurons were obtained by single cell or nucleus RNA-sequencing and then mapped to our transcriptomic taxonomy ^64^ to identify the transcriptomic type of each neuron (shown as clusters at the bottom of the dot plot). Cells are grouped by their source region, and they mainly belong to a few transcriptomic types – L2/3 IT clusters 178, 180 and 183, and L4/5 IT clusters 191, 195, 196 and 200, with some interareal difference. Within each source region, cells labeled from different projection targets (injection target region of interest, ROI) are compared, and found to be assigned to a similar subset of transcriptomic types without major distinction. cMOs or cMOp denotes contralateral MOs or MOp, respectively.

**Extended Data Figure 11.**
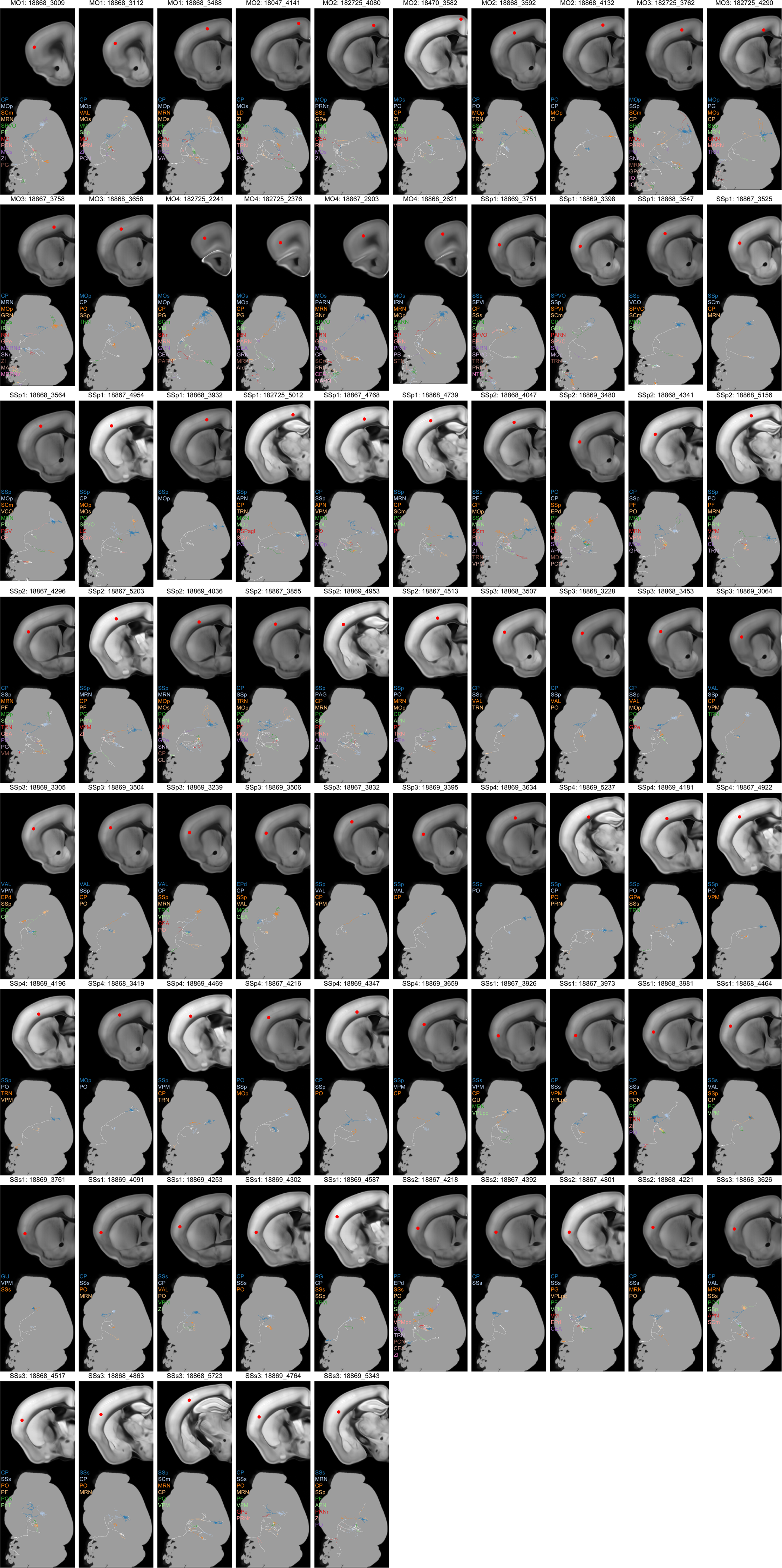
Overview of reconstructed cortical L5 ET neurons, visualized within the CCFv3 3D reference space. Each neuron is shown in two views: top, a coronal plane showing the location of the soma (red dot); bottom, a sagittal maximum projection view showing the brain-wide axon projection pattern. Cortical target regions with axon length >1 mm are indicated by different colors while other axon branches are shown in white.

**Extended Data Figure 12.**
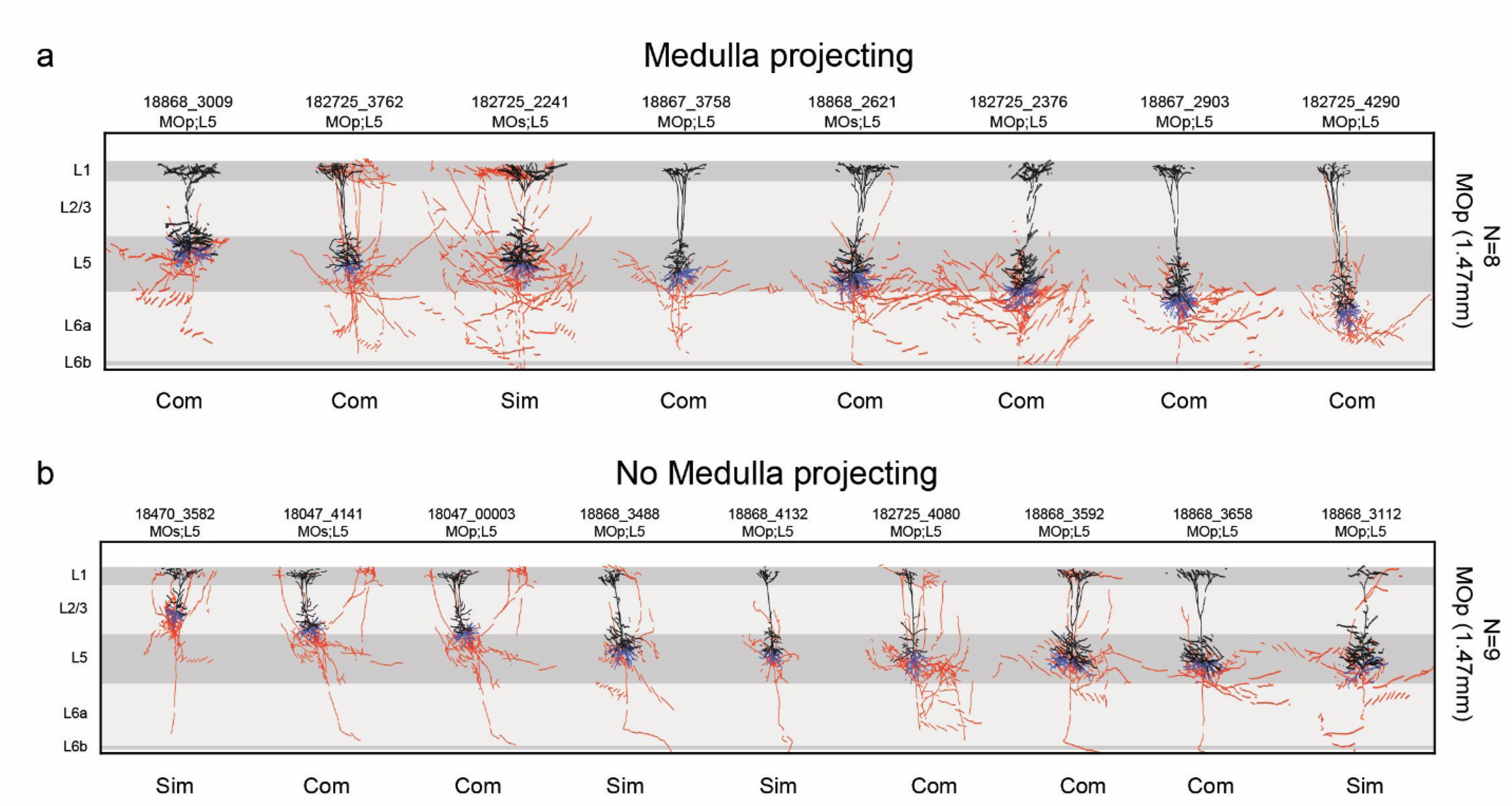
Local morphologies of motor cortex (MOp and MOs) L5 ET neurons separated into medulla-projecting and non-medulla-projecting groups. Neurons with a single apical dendrite (not branching until reaching L1) are assigned as “simple” (Sim). Neurons with two or more apical dendrites (with early branching) are assigned as “complex” (Com). Apical dendrite in black, basal dendrite in blue, axon in red. Broken lines are due to the substantially tilted nature of these MOp and MOs neurons.

**Extended Data Figure 13.**
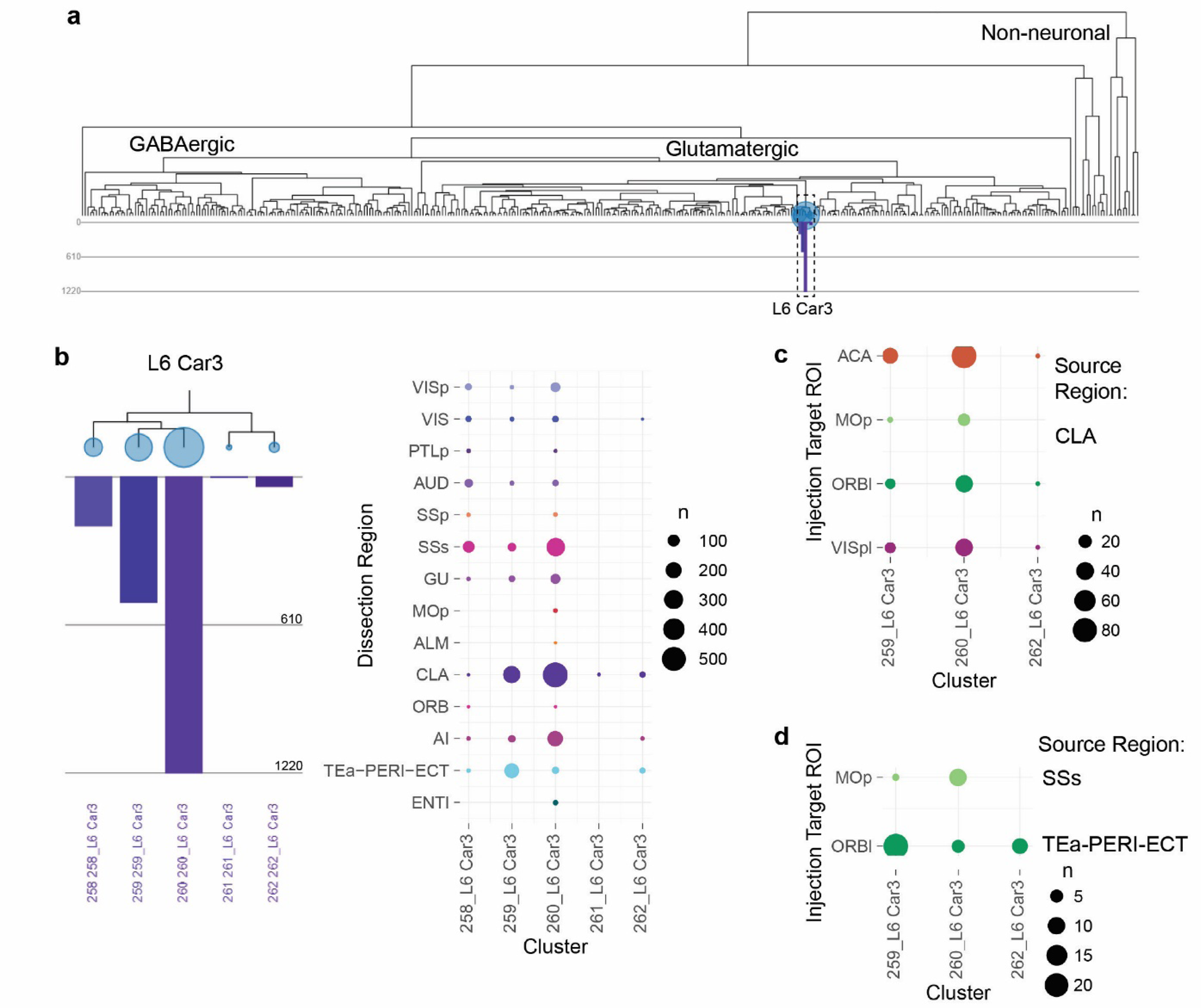
Single-cell RNA-seq characterization of L6 Car3 subclass of cortical and claustral neurons. **a,** Dendrogram of the co-clustering analysis of SMART-Seq v4 data from ∼75,000 cortical, hippocampal and claustral cells reveals a distinct branch of L6 Car3 subclass (dashed box). **b**, The L6 Car3 subclass consists of 5 clusters with variable numbers of cells in each. Dot plot (right panel) shows the number of cells from each cortical region or claustrum contributing to each cluster. Clusters with large numbers of cells are contributed by cells coming from nearly all sampled regions. **c,** Retro-seq of claustral (CLA) cells shows that CLA cells projecting to different targets (ACA, MOp, ORBl or VISpl) are mapped to the same set of transcriptomic clusters without major distinction. **d,** Retro-seq of cortical cells shows that cells from SSs projecting to MOp and cells from TEa-PERI-ECT region projecting to ORBl are mapped to a similar set of transcriptomic clusters.

**Extended Data Figure 14.**
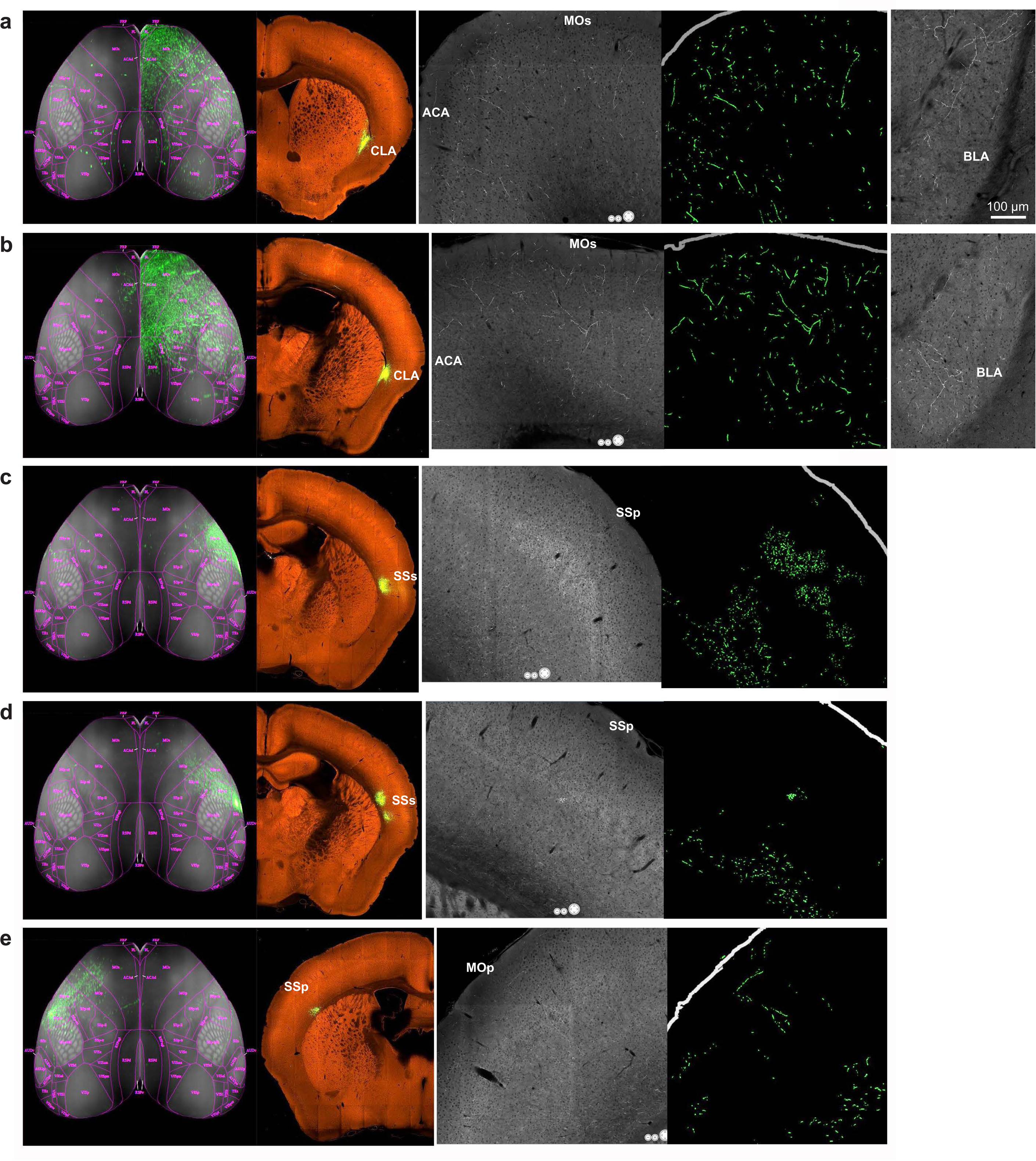
Anterograde bulk AAV tracing of projections from *Gnb4*+ neurons in claustrum or lateral cortex. **a-e,** AAV2/1-pCAG-FLEX-GFP tracer was injected into the claustrum (a-b), SSs (c-d) or SSp (e) in Gnb4-IRES2-Cre or Gnb4-IRES2-CreERT2 mice. Brains were imaged by the TissueCyte STPT system. First panel in each row: top-down view of segmented GFP-labeled axon projections in the cortex. Second panel: injection site. Third panel: the fine axon fibers in a target cortical area. Fourth panel: the segmented image of the third panel to visualize and quantify the axon fibers. Fifth panel in a and b: axon fibers observed in BLA. Full STPT image datasets are available at the Allen Mouse Brain Connectivity Atlas web portal (http://connectivity.brain-map.org/) with the following experiment IDs: a, 514505957; b, 485902743; c, 553446684; d, 581327676; e, 656688345. These 5 selected datasets all had small, spatially specific, injection sites that were located very close to each other. These small bulk injections demonstrate very distinct projection patterns between claustral and cortical *Gnb4*+ neurons.

